# Extensive distribution of β-1,2-glucanases: finding of new glycoside hydrolase families of β-1,2-glucanases

**DOI:** 10.1101/2024.02.06.578578

**Authors:** Masahiro Nakajima, Nobukiyo Tanaka, Sei Motouchi, Kaito Kobayashi, Hisaka Shimizu, Koichi Abe, Naoya Hosoyamada, Naoya Abara, Naoko Morimoto, Narumi Hiramoto, Ryosuke Nakata, Akira Takashima, Marie Hosoki, Soichiro Suzuki, Kako Shikano, Takahiro Fujimaru, Shiho Imagawa, Yukiya Kawadai, Ziyu Wang, Yoshinao Kitano, Takanori Nihira, Hiroyuki Nakai, Hayao Taguchi

## Abstract

β-1,2-Glucans are natural glucose polymers that play important physiological roles, including as symbiotic or pathogenic factors and in osmoregulation. Phylogenetically new glycoside hydrolase (GH) families have recently been identified from β-1,2-glucanase (SGL) sequences from bacteria (GH144) and a fungus (GH162). In this study, we identified four phylogenetically new groups (Groups 1–4), and determined that these families, together with GH144, GH162, and GH189, a family of transglycosylase domains in cyclic β-1,2-glucan synthases, form a superfamily. Biochemical analysis of six proteins in these groups revealed that the proteins in Groups 1–3 showed hydrolytic activity specific to β-1,2-glucan. The kinetic parameters of the enzymes of Groups 1–3 were similar to GH144 and GH162 SGLs, indicating that these enzymes were SGLs. Optical rotation analysis revealed that the SGLs followed an anomer-inverting mechanism. Structural analysis and prediction of the proteins in Groups 1–4, GH144, GH162, and GH189 suggested that Groups 1–3 and GH144 had the same reaction mechanism. Nevertheless, Groups 1–3 were dispersed irregularly in the superfamily. Overall, we determined that Groups 1–3 were new GH families, GHxxx, GHyyy, and GHzzz, respectively, and proposed that this superfamily be called an SGL superfamily because of the phylogenetical, functional, and structural relationships within the superfamily.

**Highlights:**

Wide variety of glycoside hydrolases is far beyond our understanding.
Functional and structural analysis identified three new glycoside hydrolase families.
Molecular evolution with irregular changes in reaction mechanism was revealed.

## Introduction

Carbohydrates are used not only as energy sources, but also for the storage of polysaccharides and in cell skeletons, including in the cell walls of plants and fungi [1–3]. Indigestible carbohydrates known as prebiotics affect the microbiome in the large intestine by enhancing the growth of beneficial bacteria, which is important for immuno-stimulation [4–6]. The wide variety of the physiological functions of carbohydrates in organisms derives from the complexity of the carbohydrate structures. It is estimated that over 1 billion different structures are theoretically possible for a hexasaccharide [7]. In response to such complexity, the enzymes associated with carbohydrate synthesis and degradation have evolved to have diverse amino acid sequences and tertiary structures.

Such groups of enzymes are classified into families in the Carbohydrate-Active enZYmes (CAZy) database basically based on the amino acid sequences [8–11], and are called “cazymes”. These families are categorized into glycoside hydrolase (GH) families, glycosyltransferase (GT) families, carbohydrate esterase (CE) families, polysaccharide lyase (PL) families, and auxiliary activity (AA) families (AA families contain redox-active enzymes) [9] mainly based on the types of reactions that the enzymes catalyze. Enzymes in the GH families are the most abundant in the CAZy database, and the number of identified GH families continues to increase and had reached 180 as of Sep 2024, suggesting the importance of the GH family of enzymes. However, considering the complexity of carbohydrate structures, it is presumed that many “cazymes” remain as yet unidentified.

β-1,2-Glucans are polysaccharides composed of glucose, which are found in nature mainly in cyclic forms. Although β-1,2-glucans are thought to be rare compared with the other glucans such as cellulose (β-1,4-linkage) and laminarin (β-1,3-linkage), β-1,2-glucans are involved in the bacterial infection of animal and plant cells and in hypo-osmotic adaptation [12–15]. Recently it has been reported that β-1,2-glucotriose, decomposed products of β-1,2-glucans, showed pattern-triggered immune response in plants by producing reactive oxygen species [16]. A β-1,2-glucan was first found as an extracellular polysaccharide from *Rhizobium radiobacter* (formerly *Agrobacterium tuberculosis*), a plant pathogen forming crown galls on plant roots in 1940 [17,18]. This polysaccharide was later found to be a β-1,2-glucan polymer with a cyclic form [19,20]. Enzymes for its synthesis were independently identified as cyclic β-1,2-glucan synthases (CGSs) from *Brucella abortus*, *Sinorhizobium meliloti*, and *R. radiobacter* [21–25]. The overall reaction mechanism of CGSs has been studied biochemically using a CGS from *B. abortus* [26,27].

In contrast to the enzymes for the synthesis of β-1,2-glucans, there had been no sequence-identified β-1,2-glucan-degrading enzymes until 1,2-β-oligoglucan phosphorylase (SOGP) was found in *Listeria innocua* in 2014 [28]. A method for the large-scale production of β-1,2-glucan has been established [29–31], which has enabled the investigation and identification of genes encoding other β-1,2-glucan-degrading enzymes using the produced glucans. *endo*-β-1,2-Glucanases (SGLs) that release β-1,2-glucooligosaccharides [Sop_n_s, where n is the degree of polymerization (DP)] from β-1,2-glucans have been identified from a bacterium and a fungus [32–34]. These enzymes were classified as the new GH families, GH144 and GH162 from the bacterium and the fungus, respectively [35,36]. Recently, the middle domain of CGS from *Thermoanaerobacter italicus*, a thermophilic bacterium, (TiCGS) alone was found to catalyze transglycosylation reaction. The finding of transglycosylation domain of TiCGS (TiCGS_Tg_) led to creation of GH189 [37].

Exploration of SGL and SOGP gene clusters resulted in findings of various β-1,2-glucans-associated enzymes with new functions and structures [38–43]. However, considering wide distribution of β-1,2-glucan in nature, understanding of β-1,2-glucans-associated enzymes is far from obtaining the whole picture of distribution of the related enzymes. The number of GH family of SGLs is much less than those including endo-type enzymes (endo-β-1,4-glucanases or endo-β-1,3/1,4-glucanases, 13 families; GH5–7, 10, 12, 16, 26, 44, 45, 48, 51, 74 and 124; endo-β-1,3-glucanases or β-1,3-glucanosyltransglucosylase, 12 families; GH5, 16, 17, 55, 64, 72, 81, 128, 148, 152, 157 and 158) based on the CAZy database.

GH144 SGL from *Chitinophaga pinensis* (CpSGL), a GH144-establishing enzyme, and GH162 SGL from *Talaromyces funiculosus* (TfSGL), a GH162-establishing enzyme, have only approximately 10% amino acid sequence identity [35,36]. However, CpSGL and TfSGL share a similar overall domain fold, indicating that there is a relationship between the molecular evolution of these families. GH144 and GH162 are categorized into a new GH clan, clan GH-S, based on structural classification [37]. Homology searching using CpSGL and TfSGL revealed that several groups were further found around the GH144, GH162, and GH189 families like steppingstones, and these groups, along with GH144, GH162, and GH189, were classified as a superfamily. In the present study, we characterized homologs in the newly found groups (new GH families) in the superfamily, biochemically and structurally, and found the complicated molecular evolution associated with changes in reaction mechanisms.

## Results

### Superfamily of SGLs

To understand the evolutionary relationship between GH144, GH162, and GH189, a PSI-BLAST search was performed using CpSGL (GH144), TfSGL (GH162), and TiCGS_Tg_ (GH189) as queries. Homologs with very low amino acid sequence identity with CpSGL, TfSGL, and TiCGS_TG_ were collected. Because these homologs do not belong to the GH144, GH162, or GH189 families, the PSI-BLAST search was repeated and the related homologs were identified. Then, a phylogenetic tree was constructed to investigate the phylogenetic relationships between the identified homologs. Homologs were extracted so as not to show more than a certain amino acid sequence identity with the others. The identity cut-off values were set so the number of homologs in each group did not exceed 250. Consequently, four groups were visualized around GH144, GH162, and GH189 as a phylogenetic tree and these groups were named Groups 1–4 (Fig. S1). Considering the scale in Fig. S1, these groups were clearly separated from each other. These results suggested that GH144, GH162, GH189, and Groups 1–4 form a superfamily.

GH189 is the largest group in the superfamily, and GH144 and GH162 are much larger families than are registered in CAZy database as of Sep 2024. While the GH162 proteins are distributed almost exclusively in eukaryotes, the homologs in the superfamily, except for GH162, are distributed widely in bacteria, including uncultured bacteria. Although the species were not clearly separated by the phylogenetic groups, Group 2 proteins were mainly distributed in Gram-positive bacteria. Groups 1 and 3 contained proteins from many marine bacteria and bacteria isolated from marine, sediment, and mudflat environments [44–46]. Group 1 was a smaller group than Groups 2 and 3, and Group 4 was the smallest in the proposed superfamily based on the number of collected homologs (see the Materials and Methods section for detail) [47–50]

To evaluate the reliability of the branching of the groups in the phylogenetic tree, several proteins were manually selected in each group to perform a bootstrap test using 1000 replications. Groups 1–4 were clearly separated from each other (Fig. 1), which was consistent with the fact that the amino acid sequence identities between the homologs from two different groups were very low (Fig. S2A). Most of the amino acid sequence identity values were less than 20%. The bootstrap probability of the branching points for Group 2, GH162, and at the root of Group 2 were low (6%, 45%, and 22%, respectively), and these low values implies that the order of branching among these groups was not clear.

**Figure 1.**
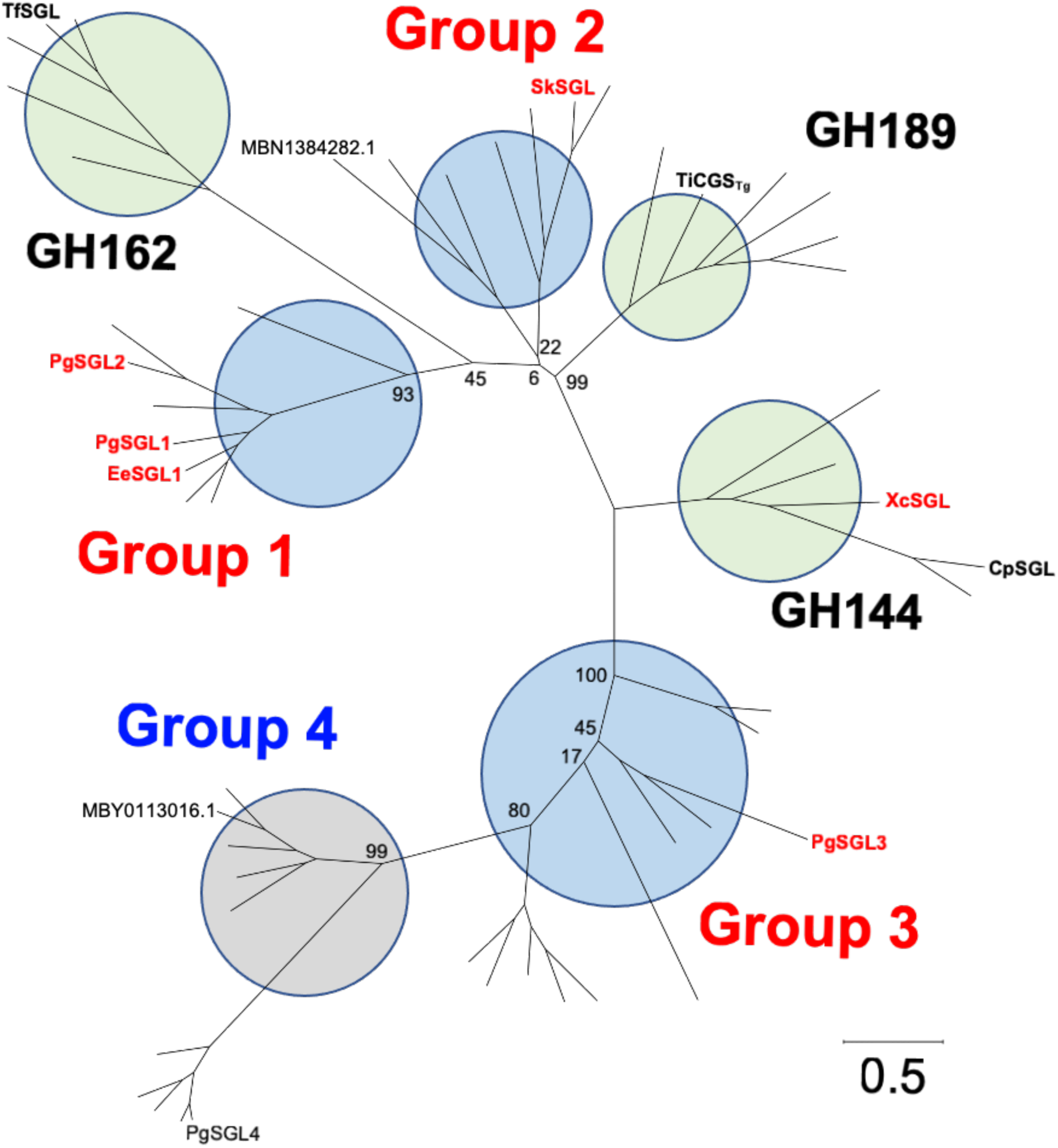
Phylogenetic tree of the superfamily of SGL-related proteins with bootstrap confidence levels. The proteins used in this study are labelled in the tree. TfSGL, CpSGL, and TiCGS_Tg_ with known biochemical functions are shown in bold black letters. The enzymes with biochemical functions that were elucidated in the present study are shown in bold red letters. Known GH families (GH144, GH162, and GH189) and newly established GH families are shown as light green and light blue circles, respectively. A group with an unknown function is indicated by the light gray circle. Bootstrap confidence levels are shown at the root branches for each group as percentages. A scale bar indicating the phylogenetic distance is shown at the bottom right. Multiple sequence alignment was performed by MUSCLE [64] and the phylogenetic tree was created using MEGA11 [65].

### General properties of the target homologs in the new groups

In this study, we used PgSGL1, PgSGL2 from *Photobacterium gaetbulicola* and EeSGL1 from *Endozoicomonas elysicola* as Group 1 proteins; SkSGL from *Sanguibacter keddieii*, a Gram-positive bacterium [51], as a Group 2 protein; PgSGL3 and PgSGL4 from *P. gaetbulicola* as Groups 3 and 4 proteins, respectively. GH144 protein from *Xanthomonas campestris* pv. *campestris* (XcSGL) was also used (see Supplementary Note, Materials and Methods, Fig. S3, Table S1 for detail). Because hydrolytic activities toward β-1,2-glucans were detected for PgSGL1, PgSGL2, PgSGL3, EeSGL1, SkSGLc and XcSGL, the general properties of these enzymes were investigated using β-1,2-glucans (NaBH_4_) (see the Materials and Methods section for details) as substrates. While many of the enzymes showed optimal pH values around neutral pH, the optimal pH values of PgSGL3 and XcSGL were 8.5 and 5.0, respectively, which suggested that the optimal pH value varied among the superfamily (Figs. S4–9). These enzymes were stable at pH ranges including neutral pH, which was consistent with the results for the optimal pH values. The optimal temperatures were 30–50 °C and the enzymes were stable at least up to 30 °C. The assay for PgSGL2 was performed in the presence of NaCl because PgSGL2 lost its activity without NaCl. The activity of PgSGL2 increased with increasing concentrations of NaCl in the reaction solution up to 500 mM (Fig. S10A). To maintain the maximum activity of PgSGL2, at least 250 mM NaCl was needed in the PgSGL2 solution (Fig. S10B). PgSGL4 did not show activity toward β-1,2-glucans nor Sop_n_s (data not shown). Size-exclusion chromatography was performed using PgSGL1, PgSGL3, and SkSGLc. PgSGL1 and PgSGL3, single domain enzymes, were eluted at the time corresponding 42 kDa, while the elution time of SkSGLc corresponded to 140 kDa (Fig. S11). These results indicated that PgSGL1 and PgSGL3 were monomeric enzymes and SkSGL was a dimeric enzyme. The *N*- and *C*-terminal domains in SkSGL may be involved in the dimer formation.

### Characteristics of the SGLs in the new groups

The substrate specificities of the six enzymes toward various polysaccharides were investigated. All the enzymes were specific for β-1,2-glucans (Table 1). The reaction patterns of these enzymes were then investigated using β-1,2-glucans as substrates. EeSGL1 (Group 1) produced Sop_3–7_ endolytically (Fig. 2A), as did PgSGL1 and PgSGL2 (data not shown). SkSGLc (Group 2) released Sop_4_ mainly in the initial phase of the reaction and produced Sop_4–8_ as the final products (Fig. 2B). Interestingly, PgSGL3 (Group 3) produced Sop_8_ predominantly in the initial stage of the reaction, and then Sop_8_ was further hydrolyzed (Fig. 2C left). When Sop_8_ was used as a substrate, Sop_8_ was hydrolyzed to Sop_3–5_ (Fig. 2C right). All the enzymes were found to be endolytic enzymes, as is CpSGL [36]Overall, in the superfamily, the substrate specificity was the same, but the degradation patterns were different. Kinetic analysis of PgSGL1, PgSGL2, PgSGL3, EeSGL1, and SkSGLc was performed (Fig. 3). All examined enzymes showed sufficiently large *V*_max_ values and sufficiently small *K*_m_ values as GH enzymes (Table 2). The five enzymes (Groups 1–3) were clearly identified as SGLs. XcSGL can be regarded as an SGL because of its narrow specificity for β-1,2-glucans and similar level of specific activity toward β-1,2-glucans as the other SGLs. Finally, the superfamily was named as a β-1,2-glucanase superfamily (SGL superfamily).

**Figure 2.**
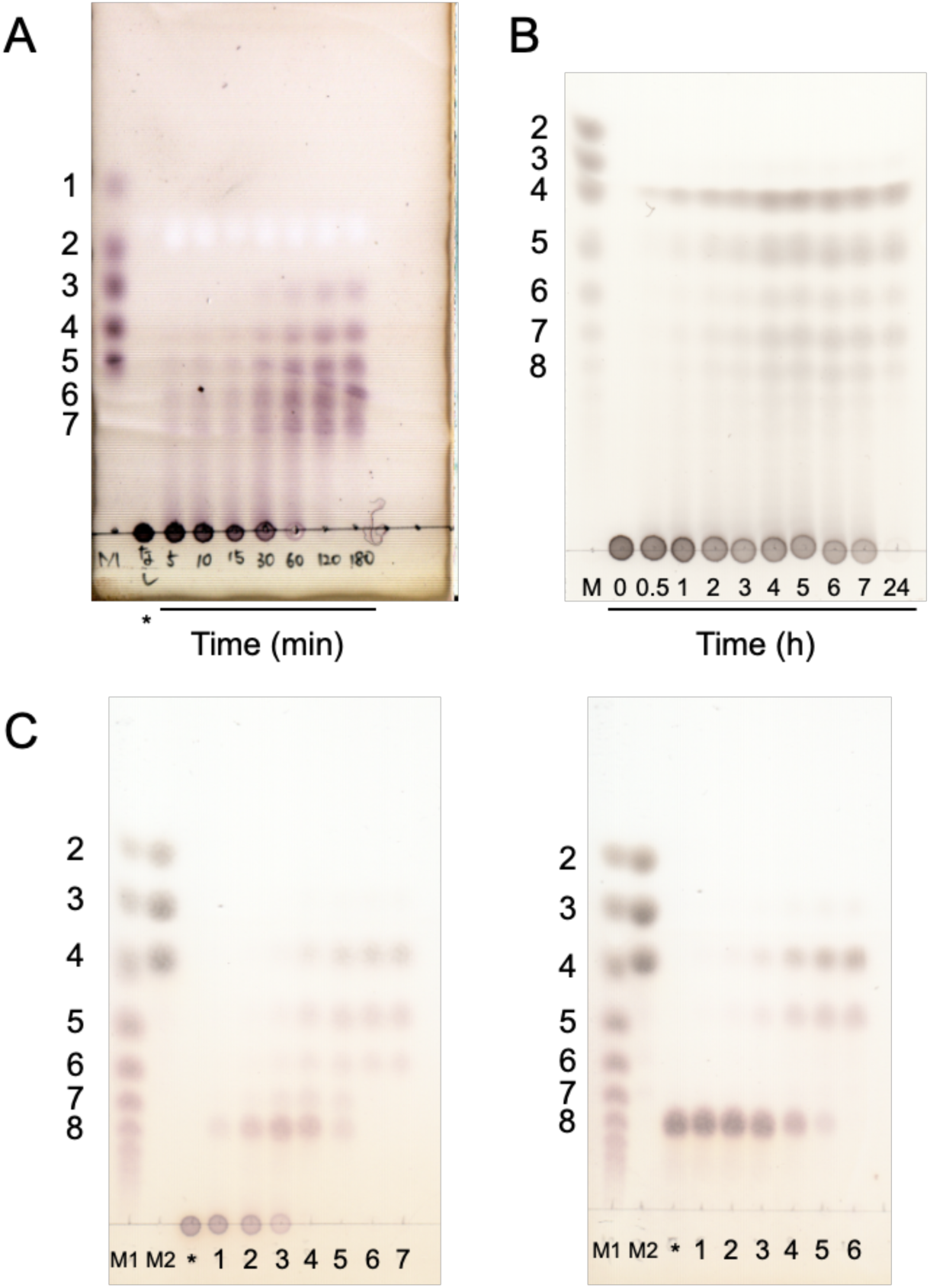
Action pattern analysis using TLC. A, EeSGL1 (Group 1); B, SkSGLc (Group 2); C, PgSGL3 (Group 3). Asterisks indicate the reaction mixture without enzymes. Lanes M, M1, and M2 show the markers. (A) The reaction times are written directly on the TLC plate. Lane M, 0.2 μl of solution containing 0.5% each Glc, Sop_2_, Sop_3_, Sop_4_, and Sop_5_. The reaction solutions (0.5 μl) were spotted on the plate. (B) Lane M, 0.5% Sop_n_s mixture prepared as described in the Materials and Methods section. A 1-μl aliquot of a marker or sample was spotted on the plate. (C) Lane M1, 1% Sop_n_s mixture prepared as described in the Materials and Methods section. Lane M2, a mixture of Sop_2_ (10 mM), Sop_3_ (7.5 mM), and Sop_4_ (5 mM) was spotted. A 0.7-μl aliquot of a marker or sample was spotted on the plate. Lanes 1–5, the reactions were performed with 0.025, 0.05, 0.1, 0.25, and 0.5 mg/ml PgSGL3, respectively, at 30 °C for 20 min. Lanes 6–7 (left), the reactions were performed with 1 mg/ml PgSGL3 at 30 °C for 20 and 220 min, respectively. Lane 6 (right), the reaction was performed with 1 mg/ml PgSGL3 at 30 °C for 30 min.

**Figure 3.**
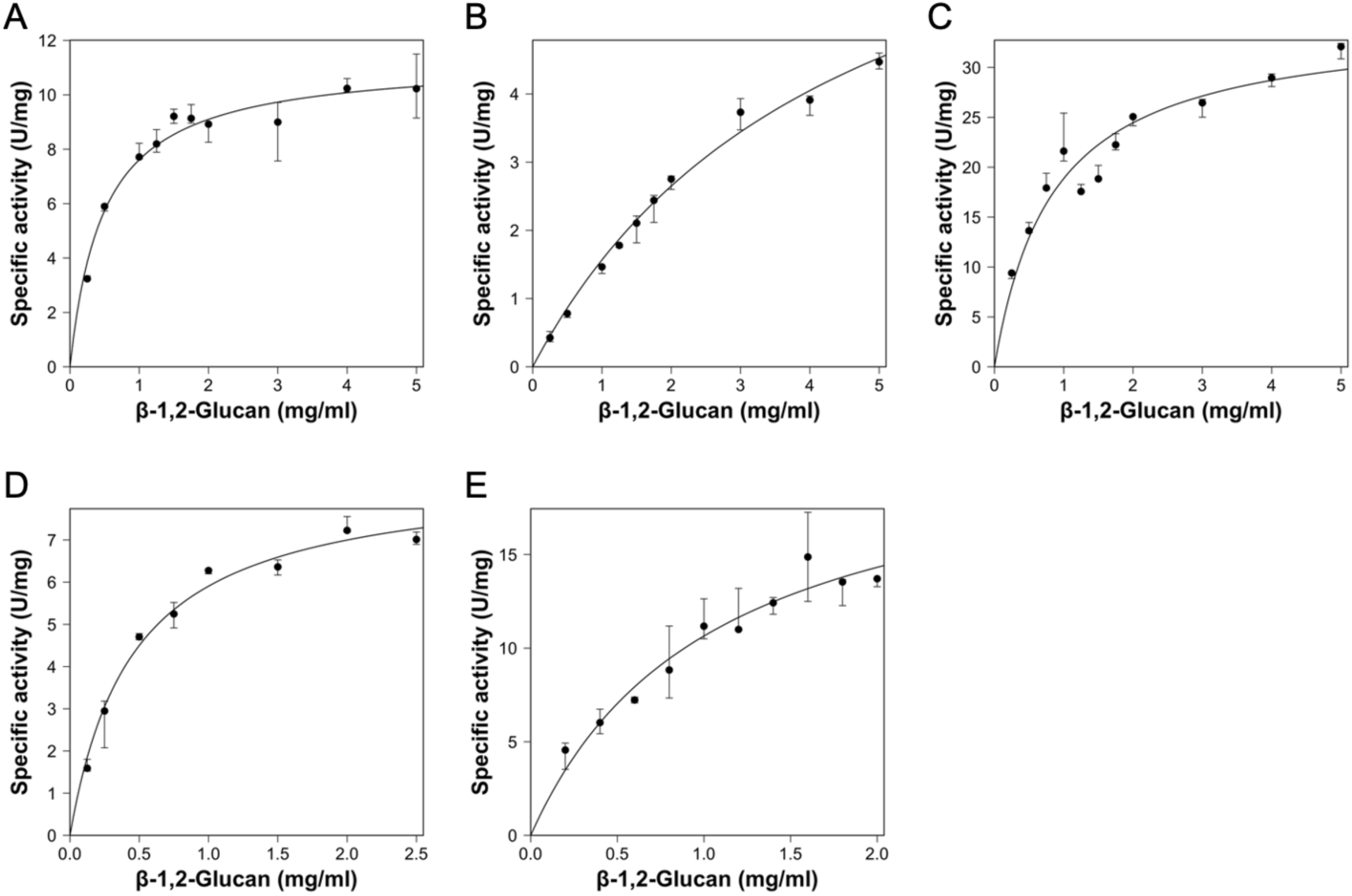
Kinetic analysis of the SGL-superfamily enzymes. Group 1, (A) PgSGL1, (B) PgSGL2, (C) EeSGL1; Group 2, (D) SkSGL; and Group 3, (E) PgSGL3. Medians are plotted as closed circles. The other data in the triplicate experiments are shown as bars. In the case of EeSGL1, two data points (1.25 and 1.5 mg/ml substrate) were eliminated from the graph because they were obviously outliers.

**Table 1.** Substrate specificity of SGL-superfamily enzymes.

| Substrate | Final<br>concentration<br>(%) | Substrate specificity (U/mg) |  |  |  |  |  |
| --- | --- | --- | --- | --- | --- | --- | --- |
|  |  | Group1 |  |  | Group2 | Group3 | GH144 |
|  |  | PgSGL1 | PgSGL2 | EeSGL1 | SkSGLc | PgSGL3 | XcSGL |
| β-1,2-Glucan<br>(NaBH<sub>4</sub>) | 0.2 | 8.44<br>(0.92)<sup>a</sup> | 3.13<br>(0.41) | 25.4<br>(3.0) | 12.3<br>(1.0) | 8.96<br>(0.85) | 3.85<br>(0.14) |
| CM-Cellulose | 0.1 | N.D.<sup>c</sup> | N.D. | N.D. | N.D. | N.D. | N.D. |
| Barley β-glucan | 0.2 | N.D. | N.D. | N.D. | N.D. | N.D. | N.D. |
| Glucomannan | 0.05 (0.1)<sup>b</sup> | N.D. | N.D. | N.D. | N.D. | N.D. | N.D. |
| Tamarind xyloglucan | 0.2 | N.D. | N.D. | N.D. | N.D. | N.D. | N.D. |
| Lichenan | 0.0125 (0.01) | N.D. | N.D. | N.D. | N.D. | N.D. | N.D. |
| Laminarin | 0.0125 (0.01) | N.D. | N.D. | N.D. | N.D. | N.D. | N.D. |
| CM-Curdlan | 0.05 | N.D. | N.D. | N.D. | N.D. | N.D. | N.D. |
| Pustulan | 0.05 | N.D. | N.D. | N.D. | N.D. | N.D. | N.D. |
| Arabinogalactan | 0.006 (0.01) | N.D. | N.D. | N.D. | N.D. | N.D. | N.D. |
| Arabinan | 0.1 (0.2) | N.D. | N.D. | N.D. | N.D. | N.D. | N.D. |
| PGA | 0.0025 (0.01) | N.D. | N.D. | N.D. | N.D. | N.D. | N.D. |
| Xylan (Beechwood) | 0.1 | N.E.<sup>d</sup> | N.D. | N.E. | N.E. | N.E. | N.E. |
| Soluble starch | 0.02 | N.D. | < 0.3% | N.D. | N.D. | N.D. | N.D. |

<sup>a</sup>The values in the parentheses after the specific activity represent the largest difference between the median value and the other data in triplicate experiments as errors.
<sup>b</sup>The values in the parentheses after the final concentrations represent the substrate concentrations for the assay of EeSGL1.
<sup>c</sup>N.D. indicates that the relative activity was lower than 0.2% of the activity toward β-1,2-glucan (NaBH<sub>4</sub>).
<sup>d</sup>N.E.: not examined.

**Table 2.** Kinetic parameters of the homologs in the SGL superfamily.

| | $V_{\max}$<br>(U/mg protein) | $K_m$<br>(mg substrate/ml) | $V_{\max}/K_m$<br>(U/mg protein<br>ml/mg substrate) |
| --- | --- | --- | --- |
| This study |  |  |  |
| (Group 1) PgSGL1 | $11.3 \pm 0.4$ | $0.484 \pm 0.068$ | $23.3 \pm 2.6$ |
| PgSGL2 | $8.58 \pm 0.71$ | $4.48 \pm 0.61$ | $1.92 \pm 0.11$ |
| EeSGL1 | $34.7 \pm 2.5$ | $0.842 \pm 0.182$ | $41.3 \pm 6.4$ |
| (Group 2) SkSGLc | $8.59 \pm 0.34$ | $0.455 \pm 0.058$ | $18.9 \pm 1.8$ |
| (Group 3) PgSGL3 | $21.8 \pm 2.5$ | $1.05 \pm 0.26$ | $20.8 \pm 2.9$ |
| References<sup>a</sup> |  |  |  |
| (GH144) CpSGL<sup>b,c</sup> | $57 \pm 4.7$ | $0.83 \pm 0.18$ | $81 \pm 12$ |
| (GH162) TfSGL<sup>b</sup> | $33 \pm 1.1$ | $0.18 \pm 0.02$ | $180 \pm 11$ |
| (GH186) OpgD from *E. coli*<sup>b</sup> | $28.7 \pm 2.0$ | $8.6 \pm 1.1$ | $3.37 \pm 0.198$ |

<sup>a</sup> The kinetic parameters of CpSGL, TfSGL, and OpgD from *E. coli* are cited from Abe et al. [36], Tanaka et al. [35] and Motouchi et al. [58], respectively.
<sup>b</sup> Kinetic parameters were evaluated using β-1,2-glucan (DP25).
<sup>c</sup> Assay was performed using the 4-hydroxybenzhydrazide method [84].

To determine the reaction mechanisms of the enzymes in Groups 1–3, i.e., anomer retaining or anomer inverting, the time course of the optical rotation during the hydrolysis of β-1,2-glucans by PgSGL1, SkSGLn (see the Experimental procedures section), and PgSGL3 was analyzed. This analysis is important to understand the fundamental characteristics that are shared in the new phylogenetic groups. The three enzymes (PgSGL1, PgSGL3, and SkSGLn) showed the same patterns as TfSGL and CpSGL that follow an anomer-inverting mechanism [35,36], indicating that these three enzymes follow an anomer-inverting mechanism (Fig. 4).

**Figure 4.**
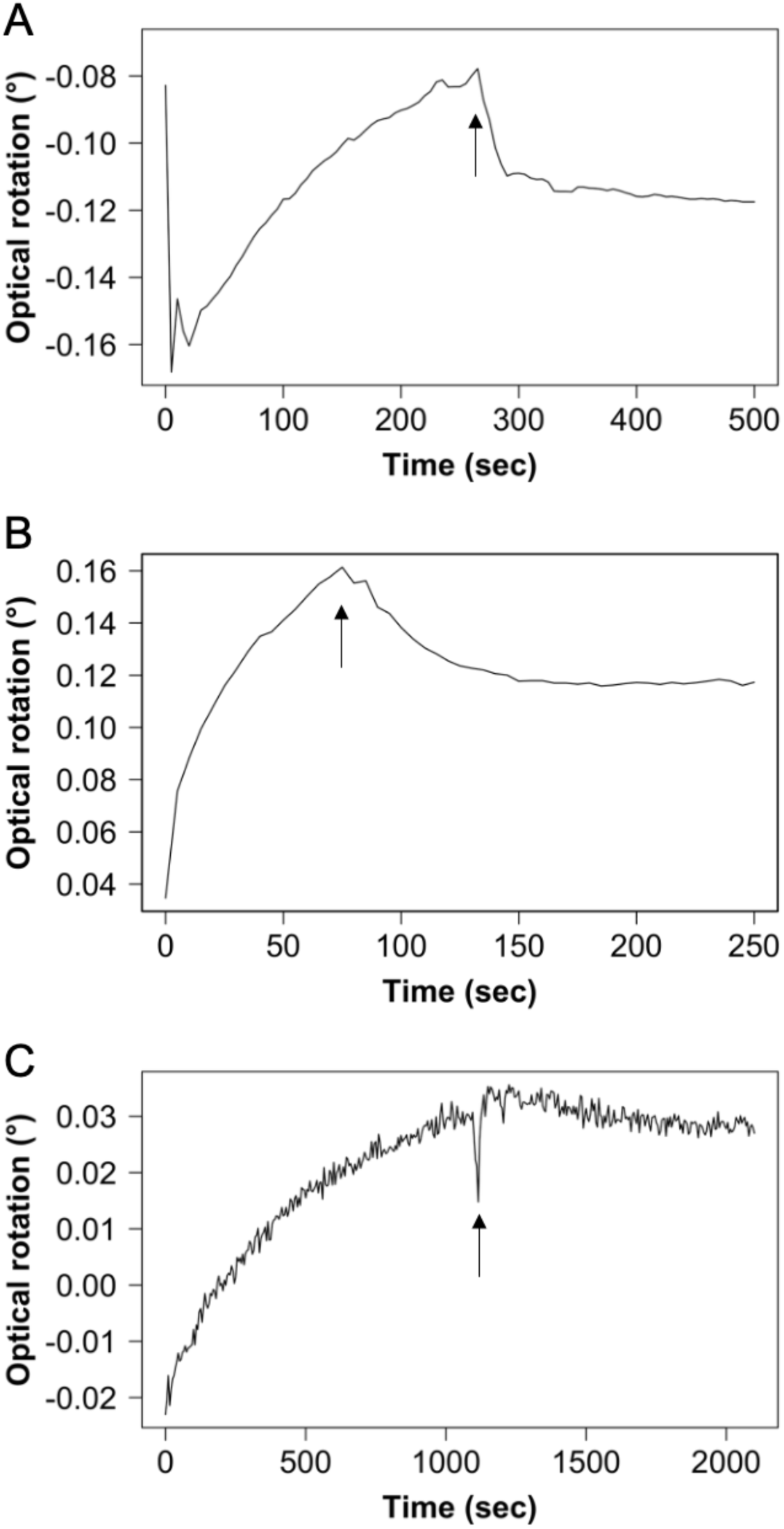
Time-course of optical rotations during the hydrolysis of β-1,2-glucans. (A) PgSGL1 (Group 1), (B) SkSGL (Group2), and (C) PgSGL3 (Group 3). The arrows indicate the time that aqueous ammonia was added to the reaction mixtures.

### Overall structural of the SGLs in the superfamily

Ligand-free structures of EeSGL1 and PgSGL3 were obtained with 2.4 and 1.2 Å resolution, respectively (Table S2). A structure of the complex with Sop_7_ was obtained with 2.5 Å resolution using XcSGL (Table S2). Overall, the structures of these enzymes are composed of a single (α/α)_6_-barrel domain, similar to CpSGL and TfSGL [35,36] (Fig. S12). SkSGL and the catalytic domain of PgSGL4 were predicted using AlphaFold2 [52]. The root mean square deviation values between the proteins of the groups in the superfamily are less than 3.5 Å according to a homology search using the DALI server (http://ekhidna2.biocenter.helsinki.fi/dali/) [53,54] (Fig. S2B), suggesting that the overall structures are similar for the proteins in the superfamily.

### Complex structures of the SGLs in the superfamily

Among the SGL superfamily, we successfully obtained complex structure of GH144 XcSGL E239Q mutant (a catalytic residue mutant) with a Sop_7_ molecule (Fig. 5A). The conformation of the Sop_7_ molecule is superimposed with that of TfSGL and both enzymes shows similar shapes of the substrate pockets (Fig, S12). Although the substrate-binding structures of EeSGL1, SkSGL, and PgSGL3 could not be determined experimentally, the shapes of the substrate pockets in these enzymes are similar to those of TfSGL and XcSGL in that the Sop_7_ molecules in TfSGL and XcSGL are accommodated in the pockets without obvious steric hindrance. Therefore, we generated substrate-binding structures computationally using molecular dynamics (MD) simulations (Fig. S13). In the final structures of the MD simulations with Sop_8_ as a substrate, the substrate is well-fitted to the pockets of EeSGL1 and PgSGL3 without appreciable structural changes. The same is true for SkSGL predicted by AlphaFold2. In the absence of a chloride ion in PgSGL3, the position of the 3-OH group of the Glc moiety at subsite +2 is deviated far from that for a putative general acid, which is inappropriate for the catalytic reaction (Fig. S14).

**Figure 5.**
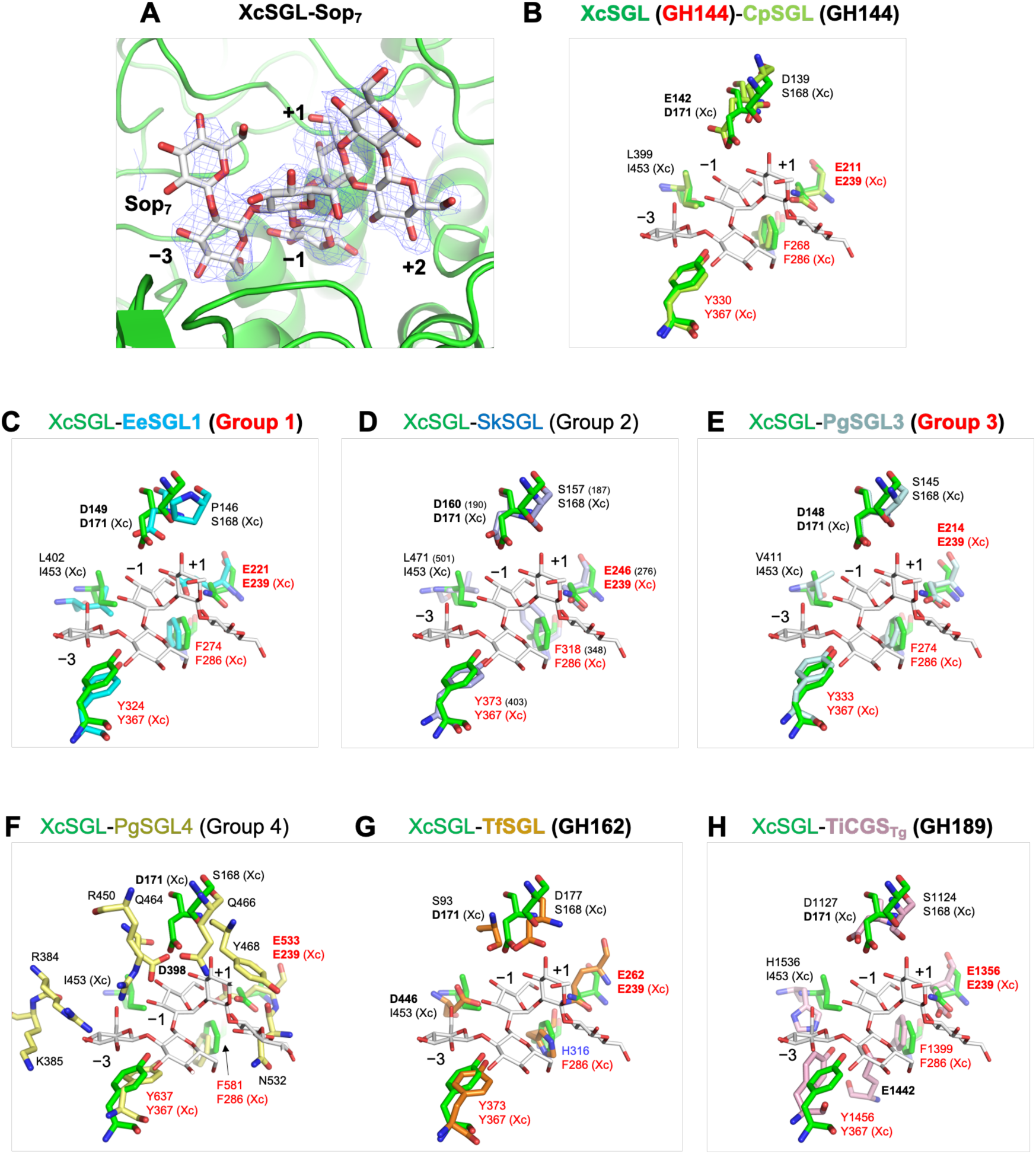
Comparison of the substrate pockets of SGLs in the SGL superfamily. (A) The complex structure of XcSGL with Sop_7_. The *F*_o_−*F*_c_ omit map for Sop_7_ is shown at the 2.0σ contour level and is represented as a blue mesh. (B–H) XcSGL (PDB ID, 8XUL) is superimposed with CpSGL (PDB ID, 5GZK) (B), EeSGL1 (PDB ID, 8XUJ) (C), SkSGL (prediction) (D), PgSGL3 (PDB ID, 8XUK) (E), PgSGL4 (prediction) (F), TfSGL (PDB ID, 6IMU) (G), and TiCGS_Tg_ (PDB ID, 8WY1) (H). Experimentally solved structures are labelled in bold, except for XcSGL in (C–H), and with bold red GH or group names. Sop_7_ molecules are represented as white sticks. The colors for each protein are as follows; XcSGL, green; CpSGL, light green; EeSGL1, cyan; SkSGL, light purple; PgSGL3, pale cyan; PgSGL4, pale yellow; TfSGL, orange; and TiCGS_Tg_, light pink. (D) The residue numbers of SkSGL are based on the starting methionine residue defined in this study. The residue numbers in parentheses are based on the sequence in the database. (F) PgSGL4 was aligned with CpSGL by pair fitting using E211, F268, and Y330 in CpSGL. (G) TfSGL was aligned with CpSGL using ligands. Sop_7_ in TfSGL is shown as a thin orange stick. The residues that define the SGL superfamily (E239, F286, and Y367 in the case of XcSGL) are colored red. The H316 in TfSGL is colored blue. Catalytic residues and candidates for catalytic residues are shown in bold letters. (Xc) represents the residues in XcSGL.

### Conservation of residues among the SGL superfamily

The conservation of residues in each group was investigated using ConSurf [55,56]. The residues forming the substrate pockets are highly conserved in each group, except Group 4 (Fig. 6), suggesting that the substrate specificity and reaction mechanisms will be the same in each group. However, the residues involved in substrate recognition are diversified across the groups in the superfamily (Fig. S15). According to the structure of PgSGL4 predicted by AlphaFold2, the substrate pocket of PgSGL4 is smaller than the pockets of the other enzymes investigated in the present study and is too small to accommodate Sop_7_, unlike in the complexes of Sop_7_ with XcSGL and TfSGL (Figs. 6E, S12). This observation is consistent with the absence of any activity of PgSGL4 toward β-1,2-glucans.

**Figure 6.**
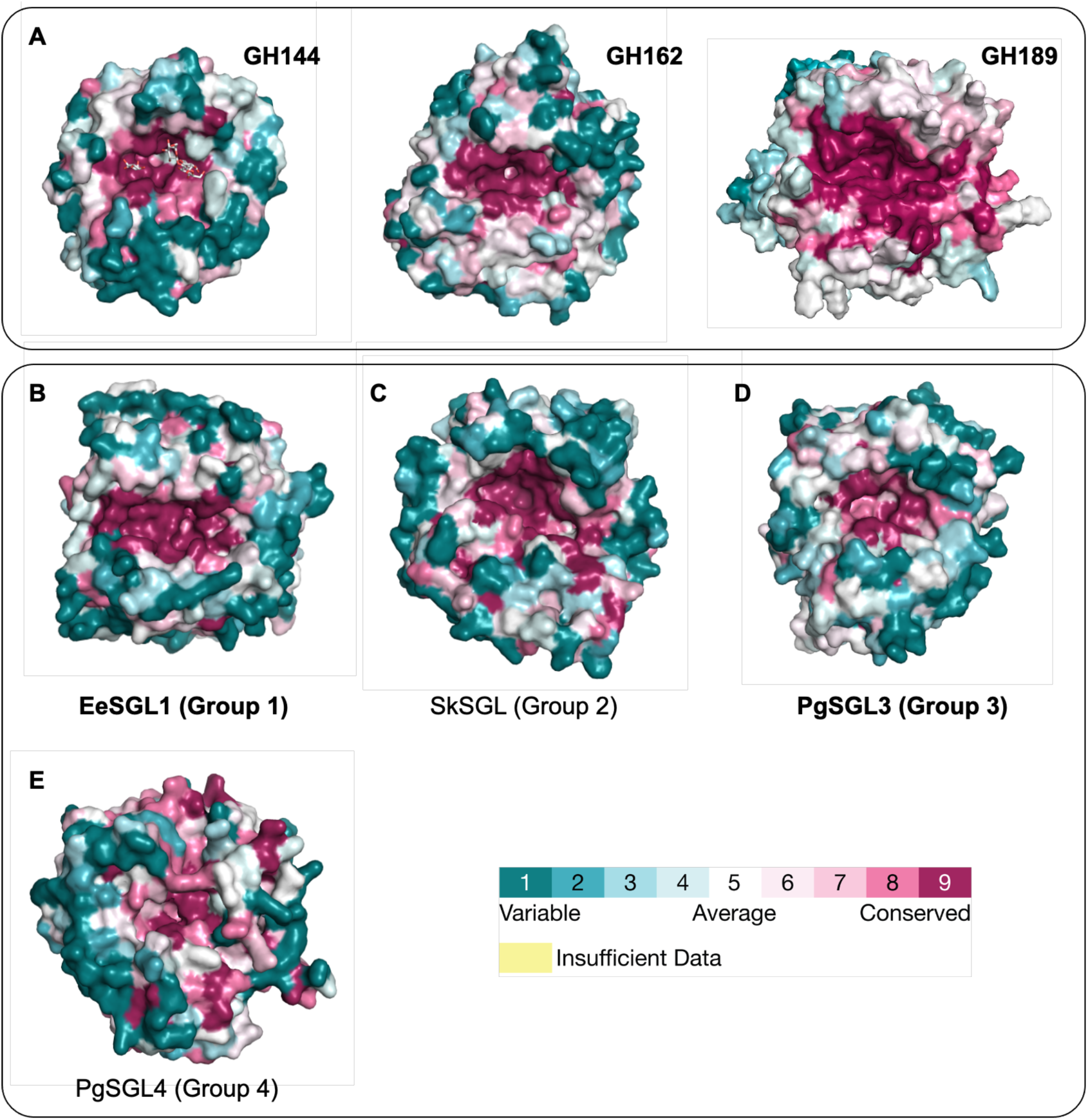
Surface models of the SGL-superfamily proteins. The SGL-superfamily proteins are shown with the surfaces colored based on the conservation scores. The conservation scores were calculated by ConSurf using the structures shown in the figure. The conservation scores are colored according to the color bar provided in the ConSurf server [55,56]. GH families or groups with experimentally solved structures and those with structures only predicted by AlphaFold2 are shown in bold and standard type, respectively. (A) GH families reported previously; (left) CpSGL (GH144, 5GZK), (middle) TfSGL (GH162; PDB ID, 6IMU), and (right) TiCGS_Tg_ (GH189, prediction). The root mean square deviation between the crystal structure of TiCGS_Tg_ (PDB ID, 8WY1) and the predicted structure was 0.54 Å. (B–E) Groups identified in this study. (B) EeSGL1 (Group 1, PDB ID, 8XUJ). (C) SkSGL (Group 2, prediction). (D) PgSGL3 (Group 3, PDB ID, 8XUK). (E) PgSGL4 (Group 4, prediction). “prediction” indicates query structures were predicted by AlphaFold2 [52].

### Mutational analysis of candidates for catalytic residues

In order to identify the catalytic residues in the newly found Groups, E221 and D149 in EeSGL1 (Group 1), and E214 and D148 in PgSGL3 (Group 3) were selected as candidate residues (see Discussion for detail). All the four mutants (E221Q and D148N in EeSGL1, E214Q and D149N in PgSGL3) showed considerably decreased hydrolytic activity toward β-1,2-glucan (less than 0.1% of the specific activity of the wild-type). This result indicated that the four residues are catalytic residues.

## Discussion

### Conserved residues defining the superfamily

A combination of multiple sequence alignments and superimposition of tertiary structures enabled the important residues shared throughout the superfamily to be identified (Figs. 5, 7 and Table S3). These residues are E239, Y367, and F286 in XcSGL (GH144) as indicated with black triangles in Fig. 7. E239 corresponds to E262 in TfSGL (GH162), which is an acid catalyst. Y367 forms a stacking interaction with a Glc moiety at subsite −3. F286 is a residue located close to a potential nucleophilic water. Although F286 is substituted for a histidine residue in GH162 (H316 in TfSGL) [35], GH162 is the only family found in eukaryotes. Thus, F286 is conserved in SGL-superfamily homologs from prokaryotes. Overall, the residues corresponding to E239, Y367, and F286 in XcSGL can be used to define the SGL-superfamily proteins. In addition, such conservation is adopted in Group 4 although its biochemical function is unknown.

**Figure 7.**
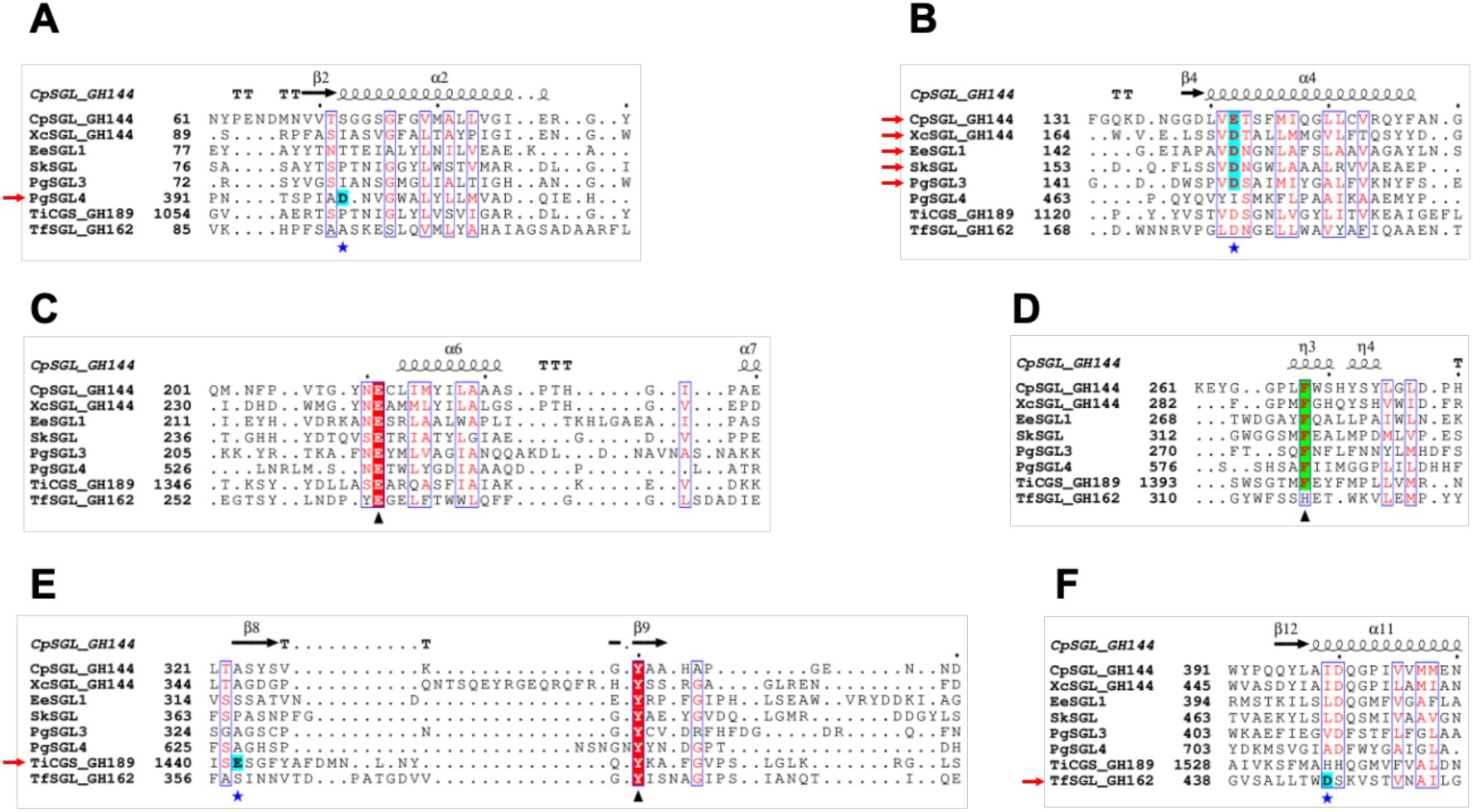
Multiple sequence alignment of the SGL-superfamily proteins. Black triangles represent highly conserved residues in the SGL superfamily, except that the phenylalanine is replaced with a histidine in GH162. The phenylalanine residues indicated by the triangle in (D) are also highlighted in green. The black triangle in (C) represents the position of a nucleophile in TiCGS_Tg_ and (candidate) base catalysts for the other proteins. Residues where blue stars and red arrows intersect are candidates for acid catalysts. These residues are also highlighted in cyan. Multiple alignment was performed using the SALIGN server [67]. The ESPript 3.0 server [68] was used for visualization of the alignment.

### The reaction mechanisms of the SGL superfamily

The similarities and differences in the reaction mechanisms between Groups 1–3 and previously reported families (GH144, GH162, GH189) are important to understand molecular evolution in the SGL superfamily. Generally, two acidic amino acids participate in reactions as catalytic residues in GH enzymes. Two catalytic residues in anomer-inverting enzymes are a general acid and a general base. A general acid residue interacts with a glycosidic bond oxygen atom directly, and a general base interacts with a nucleophilic water directly in a canonical anomer-inverting enzyme. However, the new group enzymes in the SGL superfamily are unlikely to follow the canonical mechanisms. Thus, we focus on the two catalysts for discussion below.

### General acid in the SGL superfamily

Among anomer-inverting SGLs in the SGL superfamily, the reaction route has been identified only in TfSGL (GH162) (Fig. 8A) [35]. Although the reaction route of TiCGS_Tg_ (GH189) has also been identified as shown in Fig. 8B, TiCGS_Tg_ is an anomer-retaining enzyme [37]. E262 in TfSGL (GH162), which is identified as a general acid catalyst, is conserved sequentially and spatially in the other groups in the SGL superfamily as described in the above Discussion section (Figs. 5, 7C, 8A, Table S3). This residue provides a proton to the oxygen atom in the glycosidic scissile bond through the 3-hydroxy group of the Glc moiety at subsite +2 [35] (Figs. 5G and 8A). In EeSGL1 (Group 1) and PgSGL3 (Group3), the corresponding residues are E221 and E214, respectively (Fig. 5CE), and both the E221Q mutant (EeSGL1) and E214Q mutant (PgSGL3) had considerably decreased hydrolytic activity as described above. Although mutational analysis of E246 in SkSGL is not performed, the hydrogen atom of the 3-hydroxy group of the Glc moiety at subsite +2 faces the glycosidic bond oxygen atom in SkSGL as in the case of EeSGL1, according to the MD simulations (Fig. S16), which suggests that the reaction route from E246 is appropriate for the catalytic reaction in SkSGL. Substitution of the corresponding residues in CpSGL, TfSGL, and TiCGS_Tg_ (E211Q, E262Q, and E1356Q, respectively) also drastically reduce the catalytic activity [35–37]. Thus, E221 (EeSGL1, Group 1), E246 (SkSGL, Group 2) and E214 (PgSGL3, Group 3) are considered to be general acid catalysts.

**Figure 8.**
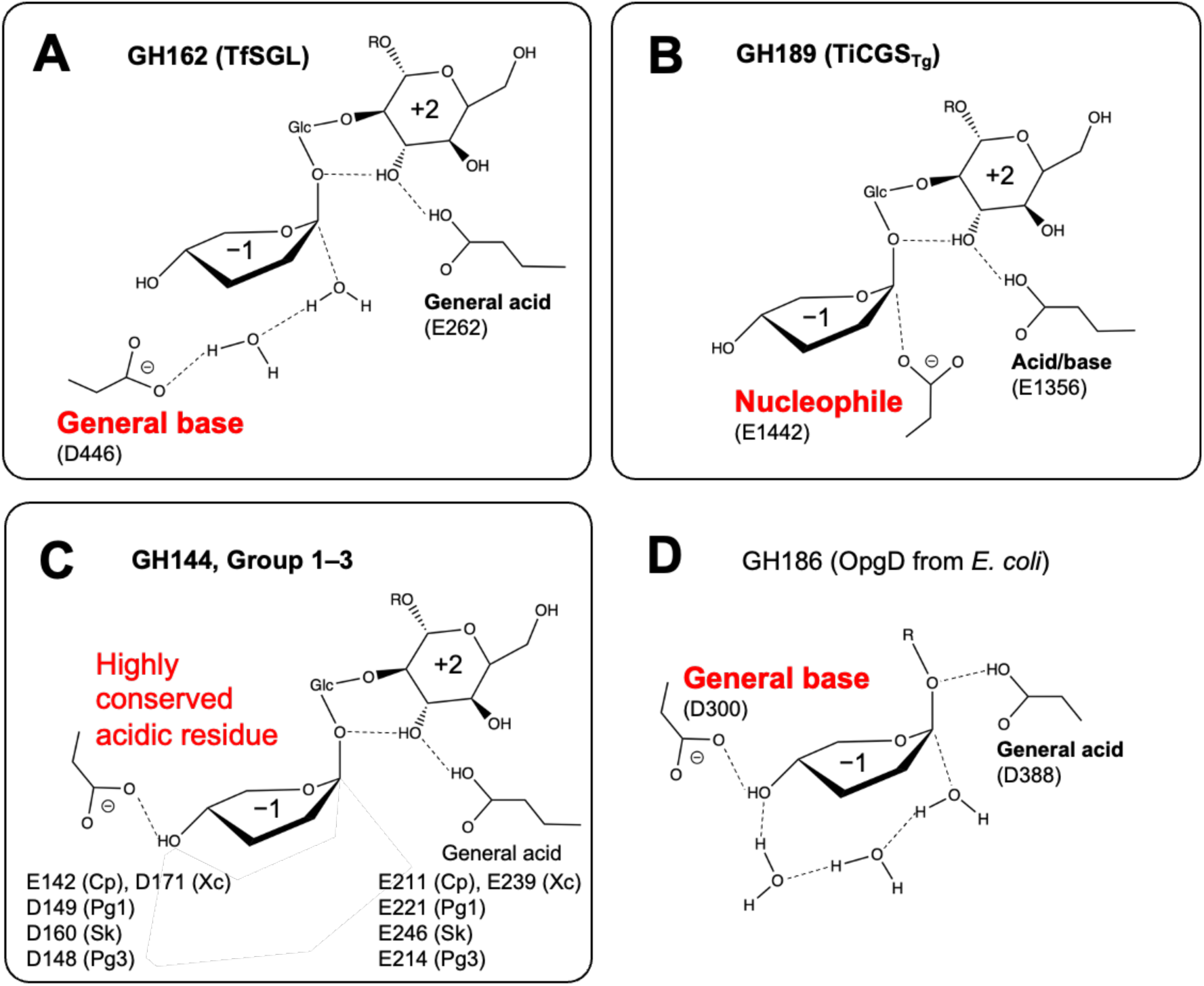
Reaction mechanisms of the SGL superfamily. Reaction mechanisms are shown schematically. (A–D) show different reaction pathways to attack the anomeric carbon at subsite −1. A nucleophile and (potential) general bases are shown in red letters to indicate the diversity in the SGL superfamily. The determined catalytic residues are shown in bold letters. Glc moieties at subsite +1 in the SGL superfamily are indicated by Glc. R represents a β-1,2- glucan moiety. Glc moieties at subsite −2 and beyond are omitted. Some substituents of the Glc moiety at subsite −1 are also omitted. (A, B) The reaction mechanisms of TiCGS_Tg_ (A) and TfSGL (B). (C) Perspective of the reaction mechanisms of GH144 and Groups 1–3. Cp, Xc, Pg1, Sk, and Pg3 are short representation of CpSGL (GH144), XcSGL (GH144), PgSGL1 (Group 1), SkSGL (Group 2) and PgSGL3 (Group 3). (D) The reaction mechanism of OpgD from *E. coli*. Groups in the SGL superfamily are bordered with solid lines (A–C).

### Diversity of general base in the SGL superfamily

In canonical anomer-inverting enzymes, a general base directly attracts a proton from a nucleophilic water molecule attacking an anomeric carbon atom at subsite −1. However, such acidic residues potentially acting on a nucleophilic water molecule are not found. Thus, candidate general base residues are discussed based on knowledge on reaction mechanisms of TfSGL (GH162) and CpSGL (GH144).

D446 acts a general base in TfSGL (GH162) and this residue corresponds to a hydrophobic residue in XcSGL and CpSGL (GH144) (Figs. 5BG, 7F), indicating that the positions of the general base catalysts are different between TfSGL and CpSGL [35,57]. However, there are several acidic amino acid residues as candidates for the general base catalyst [36]. The mutation of D135, D139, and E142 in CpSGL drastically decreases the hydrolytic activity, while the mutation of D400 retains sufficient hydrolytic activity [36], although D400 is conserved in many of the groups in the SGL superfamily (Figs. 7BF, S15). D135 is located far from subsite −1, as determined from superimposition of the structure with the XcSGL-Sop_7_ complex (Fig. 5B), and is not conserved in the SGL superfamily (Fig. 7B). D139 is also substituted for non-dissociative residues in the other groups in the superfamily. In contrast, E142 is conserved as an acidic residue in all the enzymes, except for PgSGL4 (Group 4) and TfSGL (GH162) (Fig. 7B).

The residues corresponding to E142 in CpSGL are D149 in EeSGL1 (Group 1) and D148 in PgSGL3 (Group 3) (Fig. 7B). Both the D149N mutant (EeSGL1) and D148N mutant (PgSGL3) showed drastically decreased activity as described above. Although mutational analysis of D160 in SkSGL is not performed, Sop_7_ is docked in SkSGL in almost the same conformation as EeSGL1 (Figs. 5CD, S13AB, S15). These results suggested that the residues corresponding to E142 of CpSGL are the most plausible candidates for the general base in GH144 and Groups 1–3.

Recently, GH186 SGL from *E. coli* was found to have a unique anomer-inverting reaction mechanism mediated with 4-OH group at subsite −1 and three water molecules (including a nucleophilic water) [58] (Fig. 8D). GH144 and Groups 1–3 SGLs possess acidic amino acids at the same position as the general base in GH186 SGL from *E. coli* (Fig. 8CD). This observation provides a possibility that GH144 and Groups 1–3 SGLs follow the same reaction mechanism as the GH186 SGL. However, hydrophobic environment by hydrophobic residues (I453 and F286 in the case of XcSGL, Fig. 5B–E) at the bottom of the glucose unit at subsite −1 is likely to avoid access of water molecules. Actually, no water molecule is observed at the corresponding space in the complex structure of XcSGL-Sop_7_ (Fig. 5A). These observations suggest that there could be unknown plausible reaction routes for GH144 and Group 1–3 SGLs with clearly different reaction mechanisms from that of GH162 SGL. Similarity in structural environments of catalytic pockets in GH144 and Groups 1–3 SGLs also suggests that these groups share the same reaction mechanism. In any case, acidic residues that are candidates for acid catalysts of GH144 and Groups 1–3 SGLs are located differently from those of TfSGL (GH162) and TiCSG (GH189) (Figs. 5CDEGH, 7BEF, 8ABC).

### Conclusion remarks

In this study, we found four groups with unknown functions that were related to GH144, GH162, and GH189, and identified that together these seven groups formed a superfamily. The biochemical functions were determined in three of the four newly identified groups. The three groups with identified functions are believed to share a reaction mechanism with GH144 but not with GH162 or GH189. Multiple sequence alignment revealed that only three residues were mostly conserved in the SGL superfamily. The amino acid sequence identities in the SGL superfamily were mostly less than 20% and Groups 1–4 were placed irregularly in the phylogenetic tree (Figs. 1 and S1).

An example of several GH families forming a superfamily is found with the GH94, GH149, and GH161 families of glycoside phosphorylases [59–61]. In this superfamily, the catalytic residues, inorganic phosphate recognition residues, and subsite −1 recognition residues are conserved [59], suggesting that the reaction mechanism is the same for all the members of this glycoside phosphorylase superfamily. In addition, the amino acid sequence identities in these three families are 16%–20%. Considering the criteria for classification, Groups 1–3 with functions unveiled in this study defines new GH families, GHxxx, GHyyy, and GHzzz, respectively. In addition, the overall structural similarity between the enzymes in the superfamily and the common reaction mechanism between the new families and GH144 suggest that the new GH families belong to the clan GH-S (Figs. 5, 7, S2B). The extensive distribution of SGLs found in this study suggests the importance of β-1,2-glucans and related enzymes in nature. Groups with the same reaction mechanism in the SGL superfamily are dispersed irregularly and most of substrate recognition residues are substituted between the families. These facts suggest that complicated and unique molecular evolution have occurred in this superfamily.

The identification of this superfamily showcases extensive diversity of carbohydrate-active enzymes.

## Materials and Methods

### Materials

Glucomannan, polygalacturonic acid, carboxymethyl curdlan, lichenan, arabinogalactan, barley β-glucan, tamarind xyloglucan, arabinan, and beechwood xylan were purchased from Neogen (MI, USA). Laminarin and carboxymethyl cellulose were purchased from Merck (NJ, USA). Soluble starch and pustulan were purchased from FUJIFILM Wako Pure Chemical Corporation (Osaka, Japan) and CarbioChem (CA, USA), respectively.

β-1,2-Glucans with an estimated average DP25 calculated by NMR [28] and average DP121 and DP17.7 calculated from the number average molecular weight [30,31,58] [β-1,2-Glucans (DP25), (DP121), and (DP17.7)] were prepared as reported previously. β-1,2-Glucans (DP121) used for colorimetric assay were treated with NaBH_4_ [β-1,2-glucans (NaBH_4_)] to reduce the colorization derived from the anomeric hydroxy groups at the reducing end of the polysaccharide [62].

### Sequence analysis

The amino acid sequences of TfSGL (GH162) and CpSGL (GH144) were used as queries for a PSI-BLAST search to collect 1000 and 5000 homologs, respectively. PgSGL1, PgSGL3, SkSGL, and CGSs were included in this collection. Then, a BLASTP search was performed using PgSGL1, PgSGL3, SkSGL (residues 72–541 a.a.), CGS from *B. abortus* (residues 1009–1515 a.a.; Genbank accession number, ACD71661.1), and a homolog from *Phycisphaerae bacterium* (Genbank accession number, MBN1513595.1) with the settings to collect up to 5000 homologs for the SkSGL and CGS groups and up to 1000 homologs for the other groups. Because the CGS group contained too many homologs in the database for the search, several organisms with many species were eliminated as targets for the search. The homolog from *P. bacterium* was used as a query because this homolog is located at the far side from CpSGL according to a phylogenetic tree analysis of GH144 [36]. Homologs were collected until no new group was obtained. During this collection, the PgSGL4 group, a small group, was found. The number of homolog hits found initially by BLASTP using a query in each group was approximately 5000 for GH144 CpSGL, 4100 for GH144 (MBN1513595.1), 430 for GH162 TfSGL, 1000 for SkSGL, 330 for the SkSGL group (MBN1384282.1), 480 for PgSGL3, 140 for PgSGL1, and 4200 for CGS. Because the GH144 and CGS groups were large, the GH144 proteins and GT84 domains of CGSs in the CAZy database [8,9] were also collected (1604 and 1930 homologs, respectively). Then, the duplicated sequences were removed from all the collected sequences. Multiple sequence alignment was performed against each group using Clustal Omega [63]. The transglycosylation domain and catalytic domain were extracted for GH189 and Group 4, respectively. Then, homologs showing less than 65%−95% amino acid sequence identity with the other homologs in their groups were extracted. The cut-off values were 95%, 95%, 66%, 65%, 55%, 95%, and 68% for the groups of PgSGL1, PgSGL3, SkSGL, SkSGL group (MBN1384282.1), CGS, GH162, GH144, and the homolog from *P. bacterium*, respectively. The latter two groups were combined into the GH144 group. Because the newly found PgSGL4 group was small, all the homologs in this group were used for phylogenetic analysis. Eleven homologs in the PgSGL3 group were transferred to the PgSGL4 group, and one homolog in the SkSGL group was transferred to the PgSGL1 group based on the phylogenetic positions. Finally, 20–220 homologs were obtained from each group (PgSGL1, 42; PgSGL3, 148; SkSGL, 186; PgSGL4, 20; CGS, 144; GH144, 212; and GH162, 168). All the extracted homologs containing the samples used as queries were aligned by multiple alignment using MUSCLE [64]. Then, a phylogenetic tree was constructed using the maximum-likelihood method, which was visualized by MEGA11 [65,66]. To simplify the tree, approximately five homologs, including the samples used in the present study, were taken and the tree was constructed based on the maximum-likelihood method using 1000 bootstraps. To investigate conservation of the substrate recognition and catalytic residues in the SGL superfamily, structure-based multiple sequence alignment was performed using the SALIGN server (https://modbase.compbio.ucsf.edu/salign/) [67]. This alignment was visualized using the ESPript 3.0 server (http://espript.ibcp.fr/ESPript/ESPript/) [68].

### Cloning, expression, and purification

The genomic DNA of *P. gaetbulicola* (DSM 26887), *E. elysicola* (DSM 22380), and *X. campestris* pv. *campestris* (DSM 3586) was purchased from the Leibniz Institute DSMZ (Leibniz, Germany) and *S. keddieii* ST-74 (ATCC 51767) was purchased from the American Type Culture Collection (VT, USA). Genes coding PgSGL1 (KEGG locus tag, H744_1c0224), PgSGL2 (KEGG locus tag, H744_2c01936), PgSGL3 (KEGG locus tag, H744_1c0222), PgSGL4 (KEGG locus tag, H744_1c0194), EeSGL1 (NCBI accession number, WP_026258326.1), SkSGL (KEGG locus tag, Sked_30460), and XcSGL (KEGG locus tag, XCC2207) were amplified by PCR chain reaction using KOD plus (TOYOBO, Osaka, Japan) or PrimeSTAR Max (Takara Bio, Shiga, Japan) for the DNA polymerase and the primer pairs. All the primers used in this study were designed not to include *N*-terminal signal peptides as shown in Table S4. The *N*-terminal signal peptides were predicted using SignlaP5.0 or SignalP4.1 [69,70]. The amplified PCR products were digested by the restriction enzymes shown in Table S4. Vectors (pCold I and pET30a) were also digested by the corresponding restriction enzymes suitable for ligation. The digested PCR products of PgSGL1, PgSGL2, and EeSGL1 were inserted into pCold I (Takara, Shiga, Japan) to fuse a His_6_-tag at the *N*-terminus. The digested PCR products of PgSGL3 and XcSGL, and SkSGLc were inserted into pET30a and pET24a (Merck), respectively, to fuse a His_6_-tag at the *C*-terminus. Ligation high ver. 2 (TOYOBO, Osaka, Japan) was used for the insertion.

Mutations used in this study were introduced using PrimeSTAR Max according to the manufacturer’s instructions using the primer pairs shown in Table S4. The region coding the putative *N*-terminal signal peptide of SkSGL was removed using PrimeSTAR Max and the primer pair (SkSGL Δsignal Fw and Rv). To move the *C*-terminal His_6_-tag of SkSGLc to the *N*-terminus, the gene coding SkSGLc and the pCold I vector were amplified using KOD One (TOYOBO) and the primer pairs (SkSGL pCold Fw and Rv, and pCold I Fw and Rv, respectively). The former amplified PCR product was inserted into pCold I (the latter amplified PCR product) by recombination using the SLiCE method [71]. The amplified PCR product of the gene coding PgSGL4 was inserted into pColdI by the same method as for SkSGLn. The resulting products were transferred into JM109 or XL1-blue and then the plasmids were extracted, and the target DNA sequences were checked as described above. This SkSGL fused with a His_6_-tag at the *N*-terminus was called SkSGLn.

The constructed plasmids were transferred into *E. coli* BL21(DE3). The recombinant BL21(DE3) were cultured in Luria-Bertani medium to produce target proteins. The strains with the target gene inserted into pET30a were cultured in medium containing kanamycin (30 μg/ml) at 37 °C with vigorous agitation (120 rpm) until the OD_600_ value exceed approximately 0.8 and then isopropyl β-D-1-thiogalactopyranoside (IPTG) was added to a final concentration of 0.1 mM. The cells were further cultured at 20 °C overnight. The strains with the target gene inserted into pCold I were cultured in medium containing ampicillin (100 μg/ml) at 37 °C with vigorous agitation (120 rpm) until the OD_600_ value exceeded approximately 0.6. Then, the cells were left at 10–15 °C for 30 min. IPTG was added to a final concentration of 0.1 mM and the cells were agitated again at 10–15 °C overnight. The cells were collected by centrifugation at 6000 × *g* for > 5 min and the cells were suspended in an appropriate volume of 30–50 mM MOPS (pH 7.0–7.5) or Tris-HCl (pH 7.5). The suspended cells were disrupted by sonication and the supernatants were collected by centrifugation at 33000 × *g* for > 10 min. The supernatants, except for SkSGLc, were purified by the HisTrap™ crude method (Cytiva). Each sample was loaded onto a column equilibrated with an equilibration buffer composed of 20−50 mM MOPS (pH 7.5) or Tris-HCl (pH 7.5) and 200–500 mM NaCl. The column was washed with the equilibration buffer containing 10–15 mM imidazole until almost all the unbound compounds were eluted. The samples were eluted with 50 ml of a linear gradient of up to 500 mM imidazole. For SkSGLc, other chromatography methods were used because the protein did not bind to a HisTrap™ crude column (5 ml; Cytiva, MA, USA). The cells were suspended with 50 mM Tris-HCl (pH 7.5). The supernatant was loaded onto tandemly connected HiTrap™ DEAE FF (5 ml; Cytiva) columns equilibrated with 20 mM Tris-HCl (pH 7.5). After the unbound compounds were almost all eluted with the buffer, the target protein was eluted with 100 ml of the buffer with linear gradient of 0–1 M NaCl at a flow rate of 3 ml/min. The fractions containing the target protein were collected and mixed with an equivalent volume of 20 mM Tris-HCl (pH 7.5) containing 60% saturation ammonium sulfate. After the solution was centrifugated at 33000 × *g* for 20 min, the supernatant was loaded onto a HiTrap™ Butyl HP column (5 ml, Cytiva) equilibrated with 20 mM Tris-HCl (pH 7.5) containing 30% saturation ammonium sulfate. The samples were eluted with a linear gradient of 30%–0% saturation ammonium sulfate. The fractions containing the target protein were collected. The activity in the fractionated samples was detected by TLC.

For crystallization of the XcSGL E239Q mutant (a catalytic residue mutant), the enzyme was purified by nickel affinity chromatography and further purified by hydrophobic interaction chromatography. The fractionated eluates from the HisTrapFF column were mixed with an equivalent volume of 50 mM MOPS (pH 7.0) containing 50% saturation ammonium sulfate. The sample was loaded onto a HiTrap™ Butyl column equilibrated with 50 mM MOPS (pH 7.0) containing 25% saturation ammonium sulfate. The sample was eluted with a linear gradient of 25%–0% saturation ammonium sulfate. The fractions containing the target protein was pooled and dialyzed against 5 mM MOPS (pH 7.0). For PgSGL3 and EeSGL1, the extra purification steps were not used.

The homogeneity of all the target proteins was checked using SDS-PAGE (Fig. S17). DynaMarker Protein MultiColorIII (BioDynamics Laboratory Inc., Japan) was used for protein standards. Each target protein was buffered and concentrated in 5 mM MOPS (pH 7.0) by ultrafiltration with AmiconUltra (Millipore, MA, USA) or VivaSpin (Sartorius, Germany) with molecular weight cut-off values of 10000 or 30000 at 4000 × *g*. For PgSGL2, a buffer containing 5 mM MOPS (pH 7.0) and 150 mM NaCl was used. The protein concentrations were calculated using the theoretical molecular masses and the extinction coefficients at an absorbance of 280 nm according to Pace et al. [72] and are summarized in Table S5.

### Temperature and pH profiles

To investigate the optimum pH for each enzyme, 0.2% β-1,2-glucan (NaBH_4_) was incubated with an appropriate concentration of each enzyme in 50 mM buffer (and 500 mM NaCl in the case of PgSGL2) at 30 °C. To investigate the optimal temperature, 0.2% β-1,2-glucan (NaBH_4_) was incubated with an appropriate concentration of each enzyme in the assay buffer (50 mM bicine pH 8.5 for PgSGL1; 50 mM MES, pH 6.5 containing 500 mM NaCl for PgSGL2; 50 mM MOPS pH 7.0 for EeSGL1; 50 mM cacodylate pH 6.5 for PgSGL3; 50 mM HEPES, pH 8.0 for SkSGLc; and 50 mM sodium acetate, pH 5.0 for XcSGL). To examine the pH stability, PgSGL1 (10.9 μg/ml), PgSGL2 (0.284 mg/ml), EeSGL1 (4 μg/ml), PgSGL3 (8 μg/ml), and SkSGL (5.2 μg/ml) were incubated with 50 mM buffer solutions (5 mM for EeSGL1 and PgSGL3) at 30 °C for 1 h. Then, the enzymatic reaction was performed in the presence of 0.2% β-1,2-glucan (NaBH_4_), an appropriate concentration of each enzyme, and the corresponding assay buffer. To examine the temperature stability, PgSGL1 (0.0109 mg/ml), PgSGL2 (0.284 mg/ml), EeSGL1 (4 μg/ml), PgSGL3 (8 μg/ml), and SkSGL (5.2 μg/ml) were incubated in the corresponding assay buffer (except that 5 mM MOPS pH 7.0 and 500 mM NaCl was used for PgSGL2) at various temperatures for 1 h. Then, each enzyme diluted to an appropriate concentration was incubated in the substrate solution containing 0.2% β-1,2-glucan (NaBH_4_) and the corresponding assay buffer at 30 °C. The incubation times for the enzymatic reactions were 10 min for PgSGL2, PgSGL3, and EeSGL1; 20 min for PgSGL1; and 30 min for SkSGL and XcSGL. The reaction products were colorized by the 3-methyl-2-benzo-thiazolinone hydrazone (MBTH) method described below. The assays were performed in triplicate.

### Size-exclusion chromatography

Superdex™ 200 (HiLoad 16/60; Cytiva) was equilibrated with 50 mM Tris-HCl (pH 8.0) containing 0.15 M NaCl, and each sample (0.8 mg of EeSGL1, 500 μl; 0.2 mg of PgSGL3, 500 μl) or a marker mixture (500 μl) was loaded onto the column at a flow rate of 0.5 ml/min. Ovalbumin (43 kDa), conalbumin (75 kDa), aldolase (158 kDa), ferritin (440 kDa), and thyroglobulin (669 kDa) were used as protein markers. Blue dextran 2000 was also added to the protein markers to determine the void volume of the column. The amounts of ferritin and aldolase were 0.1 and 1.5 mg, respectively, and the amounts of the other marker proteins and blue dextran 2000 were 0.5 mg. For SkSGLc, the 50 mM Tris-HCl (pH 8.0) was replaced with 20 mM Tris-HCl (pH 7.5). Elution of the samples was performed at a flow rate of 0.3 mg/ml. The molecular weights of EeSGL1, PgSGL3, and SkSGLc were calculated using equation 1,

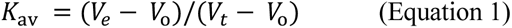

where *K*_av_ is the gel-phase distribution coefficient; *V*_e_ is the volume required to elute each protein; *V*_o_ is the volume required to elute bule dextran 2000; and *V*_t_ is the bed volume of the column.

### Substrate specificity and kinetic analysis

Enzymatic reactions were performed in the assay buffer containing each substrate at 30 °C for the times described above. The assays were performed in triplicate using the assay buffers described in the Temperature and pH profiles section. The relative activity when 0.2% β-1,2-glucan (NaBH_4_) was used as the substrate were determined for each candidate catalytic residue mutant. For kinetic analysis, the median values were plotted and fitted with a Michaelis-Menten equation (equation 2) using R.

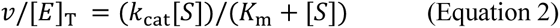

where *v* is the initial velocity; [E] is the enzyme concentration; *k*_cat_ is the turnover number; [S] is the β-1,2-glucan concentration; and *K*_m_ is the Michaelis constant.

### MBTH method

The reducing power in the reaction mixtures was determined using the MBTH method [62]. The reaction solution (20 μl), a solution containing 1 mg/ml DTT and 3.0 mg/ml MBTH (20 μl) and 0.5 M NaOH (20 μl) were mixed and then the mixture was heated at 80 °C for 30 min. After the mixture was cooled to room temperature, 40 μl of a solution containing 0.5% FeNH_4_(SO_4_)_2_·12H_2_O, 0.5% sulfamic acid and 0.25 M HCl, and 100 μl water was added. The mixtures (175 μl) were poured into a 96-well microplate (EIA/RIA plate, 96-well half area, Corning, NY, USA) and then the absorbance at 620 nm was measured. Sop_2_ was used as a standard for assaying the activity toward β-1,2-glucan (NaBH_4_) and Glc was used instead of the other substrates.

### TLC analysis

The components of the reaction mixtures of EeSGL1, PgSGL3, and SkSGLc were 50 mM MOPS (pH 7.0), 0.2% β-1,2-glucan (DP121), and 4 μg/ml of EeSGL1; 20 mM MOPS (pH 7.0), 0.2% β-1,2-glucan (DP121) or Sop_8_, and 0.025–1 mg/ml PgSGL3; and 20 mM MOPS (pH 7.5), 0.2% β-1,2-glucan (NaBH_4_), and 8.6 μg/ml of SkSGLc, respectively. The reactions were performed at 30 °C for 20 min (PgSGL3) or an appropriate time as shown in Fig. 2 (EeSGL1 and SkSGLc). The reaction was stopped by heating the sample at 80 °C for PgSGL3 or 100 °C for the other enzymes, for 5 min. The reaction mixtures and markers were spotted onto TLC Silica Gel 60 F_254_ (Merck) plates. After the plates were developed with 75% acetonitrile twice or more, the plates were soaked in a 5% (w/v) sulfuric acid/methanol solution. Spots were visualized by heating the plates in an oven. The β-1,2-glucooligosaccharide marker used for lane M1 of PgSGL3 was prepared by incubating a 1% mixture of Sop_2–5_, which are the reaction products of CpSGL, with SOGP from *L. innocua* in the presence of 1 mM sodium phosphate [58]. The Sop_n_s marker used for SkSGLc was prepared by incubating a 0.5% mixture of Sop_3–7_, which are the reaction products of SGL from *Chloroflexus aurantiacus* (Chy400_4164), with SOGP from *Enterococcus italicus* in the presence of 25 mM sodium phosphate [30,31].

### Polarimetric analysis of the reaction products

To determine the reaction mechanisms of PgSGL1, PgSGL3, and SkSGL, the time course of the degree of optical rotation in the reaction mixture was monitored. Approximately 10 ml of a substrate solution containing 2% β-1,2-glucan (DP25) [35,36] and 50 mM bicine pH 8.5 was first warmed and then poured into a cylindrical glass cell (100 mm × ø 10.5 mm). The sample was placed in a JASCO P-2200 polarimeter (Jasco, Japan) until the monitored value stabilized. Then, the reaction was started by adding PgSGL1 solution (200 μl, 40.4 mg/ml) at room temperature. When the reaction velocity began to slow down, droplets of 35% aqueous ammonia were added to enhance the mutarotation of the anomers. For PgSGL3, 2.5 mM MOPS (pH 6.5) buffer was used for the substrate solution and 300 μl of 9.6 mg/ml PgSGL3 was added. For SkSGL, 1.6 ml of substrate solution containing 1% β-1,2-glucan (DP25) and 21 mM HEPES (pH 8.0) was poured into a cylindrical glass cell (100 mm × ø 3.5 mm), because it was difficult to obtain a large amount of SkSGLn or SkSGLc and the hydrolytic activity of SkSGL was decreased in the presence of a high substrate concentration. SkSGLn solution (400 μl, 0.91 mg/ml) was added to the substrate solution.

### Crystallography

The initial screening of the crystallization conditions for the XcSGL E239Q mutant, PgSGL3, and EeSGL1 was performed using JCSG-plus™ HT-96 and PACT premier™ HT-96 (Molecular Dimensions, UK). After 1 µl of protein solution (10 mg/ml) was mixed with 1 µl of reservoir solution on a sitting plate (96-well CrystalQuick plates, Greiner Bio-One, Germany), the plate was incubated at 20 °C. The crystallization conditions were optimized from the results of the initial screening by the hanging drop vapor diffusion method using VDX plates (Hampton Research, CA, USA). Finally, crystals of the XcSGL E239Q mutant were prepared by incubation of a mixture of 11.9 mg/ml of the mutant (1 µl) and 1 µl of reservoir solution composed of 20 mM NaNO_3_, 4.5 % (w/v) PEG 3350, and 0.1 M bis-tris propane (pH 5.5) at 20 °C. For EeSGL1, the crystals obtained from the initial screening were used for data collection. The reservoir component was 0.2 M potassium thiocyanate, 0.1 M bis-tris propane (pH 6.5), and 20% (w/v) PEG 3350. For PgSGL3, a microseeding method was adopted because the crystallization was difficult to reproduce. Reservoir solution (40 mM HEPES pH 7.0, 80 mM MgCl_2_, and 10% PEG 3350) and 11.7 mg/ml of PgSGL3 (both 1 µl) were mixed and then an appropriate amount of seed crystals created by crushing a PgSGL3 crystal were added to the drop. After incubation of this hanging drop at 20 °C for 10 min, crystals were formed of suitable size for data collection.

Each crystal was soaked in the reservoir solution supplemented with 12% (w/v) PEG 200, 8% glycerol, and 4% β-1,2-glucan (DP17.7) for the XcSGL E239Q mutant, 25% PEG 400 for EeSGL1, and 25% (w/v) PEG 400 for PgSGL3 as cryoprotectants and maintained at 100 K in a nitrogen-gas stream during data collection. All X-ray diffraction data were collected on beamlines (BL-5A and NW-12A) at Photon Factory (Tsukuba, Japan). The data reduction of the diffraction data was performed using the XDS program [73]. The initial phase information was obtained by molecular replacement using the MOLREP program [74] and the structures of XcSGL and PgSGL1 predicted by AlphaFold2 [52] were used as model structures. The predicted structures were obtained from the UniProt database for the XcSGL mutant and EeSGL1. For PgSGL3, the iodide single-wavelength anomalous diffraction phasing method using the diffraction data for the iodinated PgSGL3 crystal collected at 1.8 Å was adopted [75]. The program used was the Crank2 program (shelxc, shelxd, refmac, solomon, multicomb, buccaneer, parrot) in ccp4 (http://www.ccp4.ac.uk/) [76]. Model building for EeSGL1 was performed using Buccaneer [77]. Refinement of the structures was performed using Refmac5 [78] for automatic refinement and Coot for manual refinement [79].

### MD simulations

For the MD simulations, Sop_8_ was placed in the substrate binding pockets of EeSGL1, SkSGL, and PgSGL3, referring to the complex structure of TfSGL [Protein Data Bank (PDB) ID, 6IMW]. The crystal structures of EeSGL1 and PgSGL3, and the predicted structure of SkSGL were used. To reduce calculation costs, only chain A of the EeSGL dimer and a catalytic domain of SkSGL (64–545 a.a.) were used for MD. The proteins and Sop_8_ were protonated and placed in a dodecahedral box. The box size was determined in order that all molecules were placed at least 1.5 nm from the box edges. The periodic boundary conditions were applied for all directions. The box was filled with water molecules. Sodium and chloride ions were added to each box to neutralize the total charge as well as to set the ion density to 10 mM. The AMBER ff14SB force field [80] and GLYCAM [81] were used to represent proteins and Sop_8_, respectively. The TIP3P model [82] was used for water. After energy minimization, the constant-pressure and constant-temperature (NPT) MD simulations for equilibration were performed at 1 bar and 300 K for 200 ps. Position restraints were applied to the Cα atoms of the proteins and all heavy atoms of Sop_8_ during equilibration. The production runs were performed for 100 ns at 300 K. The C-rescale method and Parrinello-Rhaman method were used to maintain the pressure during the equilibrations and the production runs, respectively. The covalent bonds of hydrogen atoms were constrained using the LINCS method, and the integration time step was 2.0 fs. MD simulations were performed by using GROMACS 2022.4 [83].

## Accession numbers

The atomic coordinates and structure factors (codes 8XUJ, 8XUK, and 8XUL) have been deposited in the PDB.

## Abbreviations

The abbreviations used are: CAZy, Carbohydrate-Active enZYmes; GH, glycoside hydrolase; GT, glycosyltransferase; CGS, cyclic β-1,2-glucan synthase; TiCGS, CGS from *T. italicus*; SOGP, 1,2-β-oligoglucan phosphorylase; SGL, β-1,2-glucanase; DP, degree of polymerization; Sop_n_, β-1,2-glucooligosaccharide (n is DP); BGL, β-glucosidase; Glc, glucose; SO-BP, a solute-binding protein in an ABC transporter specific to Sop_n_s; OPG, osmoregulated periplasmic glucans; CpSGL, SGL from *C. pinensis*; TfSGL, SGL from *T. funiculosus*; TiCGS_Tg_, transglycosylation domain of TiCGS; SkSGL, a protein from *S. keddieii* (KEGG locus tag, Sked_30460); PgSGL1, a protein from *P. gaetbulicola* (KEGG locus tag, H744_1c0224); PgSGL2, a protein from *P. gaetbulicola* (KEGG locus tag, H744_2c1936); PgSGL3, a protein from *P. gaetbulicola* (KEGG locus tag, H744_1c0222); PgSGL4, a protein from *P. gaetbulicola* (KEGG locus tag, H744_1c0194); EeSGL1, a protein from *E. elysicola* (NCBI accession number, WP_026258326.1); XcSGL, a GH144 enzyme from *X. campestris* pv. *campestris* (KEGG locus tag, XCC2207); SkSGLc, (see the Experimental procedures section for details); SkSGLn, (see the Experimental procedures section for details); MD, molecular dynamics; IPTG, isopropyl β-D-1-thiogalactopyranoside; MBTH, 3-methyl-2-benzo-thiazolinone hydrazone.

## Appendix A. Supplemental material

The following is supplementary file to the article.

Supplementary Note, Table S1–5, Figure S1–17, References

## Acknowledgments

We appreciate the help of all the staff at the Photon Factory for X-ray data collection (proposal nos. 2018G506, 2020G527, and 2022G523). We thank Edanz (https://jp.edanz.com/ac) for editing a draft of this manuscript. Yuta Takahashi, Dr. Naohisa Sugimoto, and Dr. Shinya Fushinobu supported the experiments.

## Contributions

All authors: investigation, data curation, writing – review & editing. M.N, H.T, T.N, and H.N: methodology, project administration. M.N and H.N: conceptualization. M.N: writing – original draft.

