## Supplemental for "Extensive distribution of β-1,2-glucanases: finding of new glycoside hydrolase families of β-1,2-glucanases"

Contents:

**Supplementary Note**

**Supplementary Table S1**: Classification of putative sulfatases in *P. gaetbulicola.*

**Supplementary Table S2**: Data collection and statistics.

**Supplementary Table S3**: Residues defining SGL superfamily

**Supplementary Table S4**: Primers used in the study.

**Supplementary Table S5**: Theoretical molecular masses and extinction coefficients of the recombinant proteins.

**Supplementary Figure S1**: The phylogenetic tree of the superfamily of SGL-related proteins without application of the bootstrap test

**Supplementary Figure S2**: Matrices of SGL superfamily

**Supplementary Figure S3**: A gene cluster of genes encoding PgSGL1, PgSGL3, and PgSGL4 from *P. gaetbulicola*

**Supplementary Figure S4**: pH and temperature profiles of PgSGL1 (Group 1)

**Supplementary Figure S5**: pH and temperature profiles of PgSGL2 (Group 1)

**Supplementary Figure S6**: pH and temperature profiles of EeSGL1 (Group 1)

**Supplementary Figure S7**: pH and temperature profiles of SkSGL (Group 2)

**Supplementary Figure S8**: pH and temperature profiles of PgSGL3 (Group 3)

**Supplementary Figure S9**: Optimal pH and temperature of XcSGL (GH144)

**Supplementary Figure S10**: Effect of NaCl on the hydrolytic activity of PgSGL2

**Supplementary Figure S11**: Size-exclusion chromatography analysis

**Supplementary Figure S12**: Comparison of substrate pockets between XcSGL-$Sop_7$ and TfSGL-$Sop_7$ complexes.
**Supplementary Figure S13**: Structures of the complexes obtained by MD simulations
**Supplementary Figure S14**: Structures of complexes of PgSGL3 in the presence and absence of a chloride ion
**Supplementary Figure S15**: The entire multiple sequence alignment of the SGL-superfamily proteins
**Supplementary Figure S16**: Substrate recognition by the catalytic residues
**Supplementary Figure S17**: Purification of SGL-superfamily enzymes

**References**

**Supplementary Note**

*Targets for analysis in the superfamily*

Gene clusters for potential β-1,2-glucan-associated enzymes, such as SOGPs and BGLs that prefer $Sop_n$s as substrates, were found in the genes encoding the proteins in the superfamily. Intriguingly, *Photobacterium gaetbulicola*, a bacterium isolated from mudflats [1], possesses homologs from multiple groups in the superfamily: PgSGL1 and PgSGL2 from Group 1; PgSGL3 from Group 3; and PgSGL4 from Group 4 (KEGG locus tags; H744_1c0224, H744_2c1936, H744_1c0222, and H744_1c0194, respectively) (Fig. S3).

The PgSGL4 gene forms a gene cluster with genes encoding a GH94 homolog (H744_1c0196, hereafter, H744_ is omitted), an ABC transporter homolog (1c0198–0200), GH3 homologs (1c0192, 1c0202, and 1c0206), a GH43 (subfamily 28) homolog (1c0207), and a Lac I family protein homolog (1c0197). Although there have been no reports on the characteristics of GH94 homologs in the 1c0196 subgroup, the structures and functions of SOGPs from *Listeria innocua* and *Lachnochlostridium phytofermentans* have been reported [2,3]. SOGPs are monomeric enzymes that consist of three major domains, while 1c0196 lacks an *N*-terminal domain and is assumed to form a dimer in a similar manner to other GH94 enzymes, such as cellobiose phosphorylases and chitobiose phosphorylases [4,5].

1c0192 and 1c0202 are close homologs of GH3 BGLs from *L. innocua* and *Bacteroides thetaiotaomicron* that prefer $Sop_n$s as substrates [6,7], which suggested that these GH3 homologs from *P. gaetbulicola* were $Sop_n$s-preferring BGLs. The structure of 1c0205 was the closest to the GH3 BGL from *Kluyveromyces marxianus* that has a wide substrate preference, of all the structurally available homologs [8].

GH43 is a large family mainly containing β-xylosidases and α-L-arabinofuranosidases, which is divided into 39 subfamilies [9–11]. A homolog in subfamily 28 from a metagenome has been reported to show hydrolytic activity toward arabinoxylooligosaccharides [12]. There have been no reports on the biochemical properties of close homologs of the solute-binding subunit (1c0198) in the ABC transporter. 1c0193 and 1c0203, which are sulfatase homologs, belong to the S1 group (Table S1), suggesting that these proteins are involved in the metabolism of sulfated sugars.

The PgSGL1 and PgSGL3 genes are located near the PgSGL4 gene in the *P. gaetbulicola* genome. These genes formed a gene cluster with several putative sulfatase genes (Fig. S3), although PgSGL2 gene is located independently in the genome. According to SulfAtlas (https://sulfatlas.sb-roscoff.fr/sulfatlas/index.html) [13,14], a database of sulfatases, 1c0221, 1c0223, 1c0227, and 1c0228 belong to the S1 group (Table S1). Enzymes in the S1 group are believed to be involved in the metabolism of sulfated carbohydrates to acidic sugars by acting on the sulfate groups [15,16]. 1c0214, 1c0216, and 1c0217 also belong to the S1 group, implying that the gene cluster 1c0206–1c0219 is also involved in the metabolism of sulfated carbohydrates. However, there was no gene encoding a sulfoglycosidase homolog (GH4, GH31, GH109, or GH188) in this region [17–20]. Overall, PgSGLs

1–4 are expected to be enzymes involved in β-1,2-glucan metabolism. Considering the large number of sulfatase homologs in the region (1c0192–1c0228), sulfated β-1,2-glucans might exist in nature, although to the best of our knowledge, there have been no reported examples.

*Endozoicomonas elysicola*, a bacterium isolated from the gastrointestinal tract of a mollusk, the sea slug *Elysia ornate*, possesses a PgSGL1 homolog (NCBI accession number, WP_026258326.1; EeSGL1) and the structure of EeSGL1 has been analyzed [21,22]. *E. elysicola* possesses the same gene cluster, including PgSGL3 and PgSGL4 homologs, as that of *P. gaetbulicola*.

In Group 2, the function of SGR_2427 from *Streptomyces griseus* can be speculated based on the components of the gene cluster of the *SGR_2427* gene. This gene cluster includes the gene encoding SGR_2426, a putative BGL belonging to GH1, and a solute-binding protein in an ABC transporter homologous with the SO-BP from *L. innocua*. Such gene configurations are found in various genomes, including those of *Microbacterium testaceum* StLB037 and *Nakamurella multipartita* (DSM 44233). These facts suggested that SGR_2427 is likely to be a β-1,2-glucan-associated enzyme. Unfortunately, SGR_2427 could not be obtained as a soluble protein (data not shown). *Sanguibacter keddieii*, a Gram-positive bacterium, also possesses a Group2 protein (SkSGL). An SkSGL encoding gene is located independently from the putative β-1,2-glucan-associated genes. SkSGL possesses additional domains with unknown functions at the *N*- and *C*-termini. A GH144 homolog from *Parabacteroides distasonis*, a common bacterium in the large intestine [23], with unknown-function domains at the *N*-terminus, was found to be a novel exo-type $Sop_2$-releasing enzyme. The domains played a critical role in the exolytic degradation activity of the enzyme[24]. Therefore, SkSGL was used as a target for Group 2.

A Michaelis complex structure has been reported for GH162 but not for GH144. To compare the functions, structures, and reaction mechanisms of the homologs characterized in the present study, a GH144 enzyme from *Xanthomonas campestris* pv. *campestris*, a phytopathogen [25,26] (KEGG locus tag, XCC2207; XcSGL), was used to obtain a Michaelis complex with β-1,2-glucans.

**Table S1. Classification of putative sulfatases in *P. gaetbulicola*.**

| Family / Subfamily[a] | UniProt accession No. | Locus |
|---|---|---|
| In the gene cluster | | |
| S1_8 | A0A0C5WJJ0 | H744_1c0228 |
| S1_11 | A0A0C5WJC1 | H744_1c0203 |
| S1_11 | A0A0C5WJA2 | H744_1c0193 |
| S1_11 | A0A0C5WGE1 | H744_1c0221 |
| S1_11 | A0A0C5WJI5 | H744_1c0223 |
| S1_11 | A0A0C5WDX6 | H744_1c0227 |
| S1_19 | A0A0C5WQP8 | H744_1c0214 |
| S1_19 | A0A0C5WDX0 | H744_1c0217 |
| S1_19 | A0A0C5WGD5 | H744_1c0216 |
| none | A0A0C5W1R0 | H744_1c0215 |
| The others | | |
| S1_9 | A0A0C5WQ05 | H744_2c1766 |
| S1_13 | A0A0C5WB34 | H744_2c2132 |
| S1_13 | A0A0C5X0G6 | H744_2c2131 |
| S1_13 | A0A0C5WTU2 | H744_1c1448 |
| S1_26 | A0A0C5WSF7 | H744_1c0940 |
| S1_27 | A0A0C5WTP4 | H744_2c1768 |
| S1_44 | A0A0C5WQ21 | H744_2c1791 |
| S3 | A0A0C5W3A8 | H744_1c0866 |
| S3 | A0A0C5WU63 | H744_1c1582 |
| none | A0A0C5W3F7 | H744_1c0941 |
| none | A0A0C5WJS1 | H744_1c1445 |
| none | A0A0C5WA56 | H744_2c1790 |
| none | A0A0C5WUR0 | H744_2c2120 |
| none | A0A0C5WVI3 | H744_2c2416 |

[a] Classification of the family was based on SulfAtlas [13]

**Table S2. Data collection and statistics.**

| Data set | EeSGL1 | PgSGL3 | XcSGL (E239Q)-$Sop_7$ |
|---|---|---|---|
| **Data collection** | | | |
| Beamline | KEK BL-5A | KEK NW-12A | KEK BL-5A |
| Space group | $P2_12_12$ | $P2_12_12_1$ | $P4_332$ |
| | $a$ = 110.50 | $a$ = 52.28 | |
| Unit cell parameters (Å) | $b$ = 114.91 | $b$ = 76.60 | $a = b = c$ = 220.18 |
| | $c$ = 78.89 | $c$ = 94.43 | |
| Resolution (Å)[a] | 46.44–2.40 (2.49–2.40) | 47.21–1.20 (1.22–1.20) | 49.23–2.50 (2.56–2.50) |
| Total reflections[a] | 529415 (57393) | 732055 (35275) | 2469753 (176787) |
| Unique reflections[a] | 40027 (4131) | 114009 (5320) | 63337 (4390) |
| Completeness (%)[a] | 100 (100) | 96.0 (91.5) | 100 (100) |
| Multiplicity[a] | 13.2 (13.0) | 6.4 (6.6) | 39.0 (40.3) |
| Mean $I/\sigma$ ($I$)[a] | 14.1 (2.9) | 11.6 (4.0) | 35.4 (6.0) |
| $R_{merge}$ (%)[a] | 14.1(93.2) | 9.6 (43.3) | 11.5 (87.6) |
| $R_{pim}$ (%)[a] | 5.7 (37.2) | 6.0 (27.1) | 2.6 (19.6) |
| $CC_{1/2}$[a] | (0.888) | (0.904) | (0.961) |
| **Refinement** | | | |
| Resolution (Å) | 46.442–2.40 | 47.213–1.200 | 49.233–2.50 |
| No. of reflections | 37877 | 108263 | 60057 |
| No. of atoms | 6738 | 3819 | 3720 |
| No. of water molecules | 9 | 332 | 22 |
| $R_{work}/R_{free}$ (%) | 22.4/28.1 | 19.1/20.8 | 33.8/36.4[b] |
| No. of asymmetric units | 2 | 1 | 1 |
| RMSD from ideal values | | | |
| Bond lengths (Å) | 0.0053 | 0.0130 | 0.0590 |
| Bond angles (°) | 1.4197 | 1.9403 | 2.0065 |
| Average $B$-factors (Å$^2$) | | | |
| Protein (chain A/B) | 46.2/45.4 | 11.9 | 41.9 |
| Ligand | | | |
| $Sop_7$ | - | - | 50.9 |
| Solvent | 34.4 | 19.6 | 33.0 |
| Ramachandran plot (%) | | | |
| Favored | 95.5 | 98.1 | 95.0 |
| Allowed | 4.5 | 1.9 | 4.5 |
| Outlier | 0.0 | 0.0 | 0.5 |
| **PDB entry** | 8XUJ | 8XUK | 8XUL |

[a] Values in parentheses are for outer shells.

[b] $R$ values are large because of the disorder of the $C$-terminal region compared with the catalytic domain. However, the assignment of XcSGL molecules was conducted using only the catalytic domain.

**Table S3. Residues defining SGL superfamily**

| Group | Enzyme | Phe | Tyr | Glu |
|---|---|---|---|---|
| Group 1 | EeSGL1 | F274 | Y324 | E221 |
| Group 2 | SkSGL | F318 (348)[a] | Y373 (403) | E246 (276) |
| Group 3 | PgSGL3 | F274 | Y333 | E214 |
| Group 4 | PgSGL4 | F241 | Y297 | E193 |
| GH144 | XcSGL | F286 | Y367 | E239 |
| GH144 | CpSGL | F268 | Y330 | E211 |
| GH162 | TfSGL | H316* | Y373 | E262 |
| GH189 | TiCGS$_{Tg}$ | F1399 | Y1456 | E1356 |

[a] The residue numbers in parentheses are based on the sequence in the database.

* Asterisk represents that a residue type is not conserved.

**Table S4. Primers used in the study.**

| Name | Sequence (5′ to 3′)[a] | Restriction enzyme |
|---|---|---|
| Cloning | | |
| PgSGL1 Fw | GTATCTCGAGCCACATAACATCCAAGTTC | XhoI |
| PgSGL1 Rv | CGGCGAATTCTTATTCGTTATTCGGAGTAAAC | EcoRI |
| PgSGL2 Fw | GATACTCGAGCTTGGTGGATGTGCATCGAGTG | XhoI |
| PgSGL2 Rv | GCCGGAATTCCTACGAAGCGTTTTCCTCACG | EcoRI |
| PgSGL3 Fw | GATGGCACATATGCAAACGGCGGCAAATG | NdeI |
| PgSGL3 Rv | GACGTCTCGAGCTCAAAGTAGTTGTTG | XhoI |
| EeSGL1 Fw | GTATCTCGAGAGCGCTGTCACACTGGATAAACTG | XhoI |
| EeSGL1 Rv | GCCGGAATTCTTACTTAGCCCGGTCGTCTTTAAAAC | EcoRI |
| PgSGL4 Fw | TATCGAAGGTAGGCATATGTGTGGTGGTGGCTCTGGC | |
| PgSGL4 Rv | TTTAAGCAGAGATTACCTAGTAGCACTGAGCACTATTATTAC | |
| XcSGL Fw | CTGCAAACATATGGAGGAGCCCAAG | NdeI |
| XcSGL Rv | AAGGCGCACCTCGAGCTCGGGTTTGGG | XhoI |
| SkSGL Fw | AAAAAAACATATGGGATCCCCCACCCGCGG | NdeI |
| SkSGL Rv | AATCGTGGCGGCCGCGTCCCGGCGACGCGTCCGGA | NotI |
| SkSGL Δsignal Fw | ACATATGGCGCCCACGCCCGAGCAG | |
| SkSGL Δsignal Rv | GTGGGCGCCATATGTATATCTCCTTC | |
| SkSGL pCold Fw | TATCGAAGGTAGGCATATGGCGCCCACGCCCGAGCAGACA | |
| SkSGL pCold Rv | TTTAAGCAGAGATTACCTAGTCCCGGCGACGCGTCCGGA | |
| pCold I Fw | TAGGTAATCTCTGCTTAAAAGCAC | |
| pCold I Rv | CATATGCCTACCTTCGATATGATG | |
| Mutants | | |
| PgSGL3 E214Q Fw | TTCAATCAATATATGCTTGTTGCCGGT | |
| PgSGL3 E214Q Rv | CATATATTGATTGAACGCTTTGGTACG | |
| XcSGL E239Q Fw | TACAACCAAGCCATGATGCTGTACATC | |
| XcSGL E239Q Rv | CATGGCTTGGTTGTAGCCCATCCAGTC | |
| EeSGL1 D149N Fw | GCTGTTAACAATGGCAACCTGGCGTTT | |
| EeSGL1 D149N Rv | GCCATTGTTAACAGCTGGCGCAATTTC | |
| EeSGL1 E221Q Fw | GCCAATCAAAGCCGACTGGCGGCACTC | |
| EeSGL1 E221Q Rv | TCGGCTTTGATTGGCCTTGCGATCCAC | |
| PgSGL3 D148N Fw | CCGGTTAACAGTGCCATCATGATTTAT | |
| PgSGL3 D148N Rv | GGCACTGTTAACCGGACTCCAGTCATC | |
| PgSGL3 E214Q Fw | TTCAATCAATATATGCTTGTTGCCGGT | |
| PgSGL3 E214Q Rv | CATATATTGATTGAACGCTTTGGTACG | |

[a] Restriction sites for NdeI and XhoI are underlined.

**Table S5. Theoretical molecular masses and extinction coefficients of the recombinant proteins.**

| | Theoretical molecular mass (g/mol) | Extinction coefficient ($A_{280}$ $cm^{-1}$ $M^{-1}$) | N-terminal signal peptide |
|---|---|---|---|
| PgSGL1 | 51432.45 | 72810 | None |
| PgSGL2 | 53698.186 | 89395 | None |
| EeSGL1 | 50508.48 | 99240 | 1–22 a.a.[a] |
| PgSGL3 | 48951.656 | 91010 | 1–21 a.a. |
| SkSGLc[b] | 73667.049 | 118190 | 1–32 (59) a.a.[c] |
| SkSGLn[b] | 77634.668 | 118315 | 1–32 (59) a.a.[c] |
| XcSGL | 60466.116 | 128230 | 1–27 a.a. |
| PgSGL4 | 126166.348 | 164155 | 1-24 a.a. |

[a] SignalP4.1 was used to predict an *N*-terminal signal peptide.

[b] Because of a sequence error (deletion of the cytosine at position 123) in the database, the corrected sequence was used.

[c] The signal region was obscure when the first methionine was adopted as a start codon. Thus, the second methionine was adopted instead. The numbering starts at the second methionine. The numbers in parentheses indicate the residue numbers when the first methionine was adopted as a start codon.

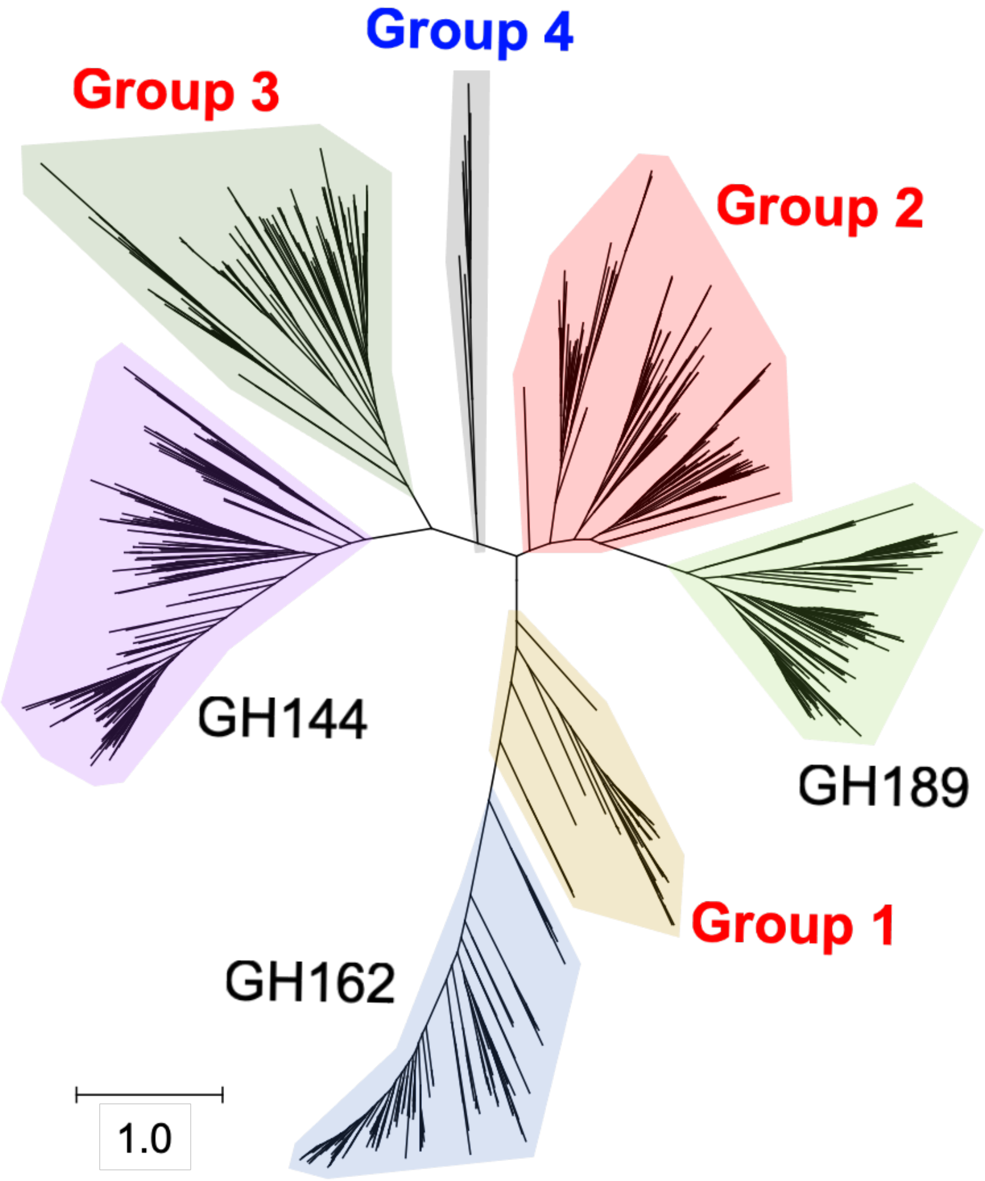

**Figure S1. The phylogenetic tree of the superfamily of SGL-related proteins without application of a bootstrap test**

Up to 250 proteins were collected from each group to create the phylogenetic tree using the method described in the Sequence Analysis section. Biochemically identified groups are labelled in bold red letters (Groups 1–3) and the unidentified group is in bold black letters (Group 4). Groups that have already been defined are labelled with the family numbers (GH144, GH162, and GH189). The groups are colored light purple (GH144), light blue (GH162), light green (GH189), light yellow (Group 1), light red (Group 2), dark green (Group 3), and light gray (Group 4). A scale bar is provided in the bottom left corner to show the phylogenetic distances.

**A**

| | | GH144 | | GH162 | GH189 | Group 1 | | Group 2 | Group 3 | Group 4 | | The number of homologs used in this study | Identity cut-off (%) |
|---|---|---|---|---|---|---|---|---|---|---|---|---|---|
| | | Cp | Xc | Tf | Ti | Pg1 | Ee1 | Sk | Pg3 | Pg4 | MBY | | |
| GH144 | CpSGL | | | | | | | | | | | 212 | 68 |
| | XcSGL | 28.3 | | | | | | | | | | | |
| GH162 | TfSGL | 11.9 | 14.0 | | | | | | | | | 168 | 95 |
| GH189 | TiCGS$_{Tg}$ | 17.6 | 15.0 | 12.7 | | | | | | | | 144 | 65 |
| Group 1 | PgSGL1 | 13.6 | 19.2 | 13.4 | 17.5 | | | | | | | 140 | 95 |
| | EeSGL1 | 15.0 | 17.1 | 13.0 | 19.9 | 52.5 | | | | | | | |
| Group 2 | SkSGL | 17.5 | 19.8 | 11.7 | 25.9 | 19.5 | 21.5 | | | | | 186 | 66 |
| Group 3 | PgSGL3 | 14.9 | 17.1 | 11.4 | 14.0 | 15.7 | 13.1 | 10.7 | | | | 148 | 95 |
| Group 4 | PgSGL4 | 12.1 | 11.1 | 11.1 | 12.0 | 14.1 | 10.0 | 12.0 | 17.4 | | | 20 | none |
| | MBY0113016.1 | 15.4 | 14.6 | 13.5 | 13.7 | 15.4 | 13.7 | 13.4 | 16.9 | 24.2 | | | |

**B**

| | | | GH144 | | GH162 | GH189 | Group 1 | Group 2 | Group 3 | Group 4 |
|---|---|---|---|---|---|---|---|---|---|---|
| | | | Cp | Xc | Tf | Ti | Ee1 | Sk | Pg3 | Pg4 |
| GH144 | CpSGL | 5GZK_A | | | | | | | | |
| | XcSGL | 8XUL | 1.6 | | | | | | | |
| GH162 | TfSGL | 6IMU_A | 2.6 | 2.7 | | | | | | |
| GH189 | TiCGS$_{Tg}$ | 8WY1_A | 2.3 | 2.3 | 2.7 | | | | | |
| Group 1 | EeSGL1 | 8XUK_A | 2.4 | 2.4 | 2.3 | 2.3 | | | | |
| Group 2 | SkSGL | AF2 | 2.4 | 2.1 | 2.5 | 1.8 | 2.0 | | | |
| Group 3 | PgSGL3 | 8XUJ | 3.0 | 2.5 | 3.1 | 2.5 | 2.9 | 2.5 | | |
| Group 4 | PgSGL4 | AF2 | 3.0 | 3.0 | 3.4 | 3.1 | 3.3 | 2.9 | 3.2 | |

**Figure S2. Matrices of SGL superfamily**

The enzymes in Groups 1–3 are shown in bold red letters and are highlighted in light blue. XcSGL, a biochemically identified enzyme in this study is highlighted in light green. Shortened names are used as column names for the superfamily proteins. Only the $(\alpha/\alpha)_6$-barrel domain regions were used; XcSGL, residues 72–498 a.a.; SkSGL, residues 42–514 a.a.; PgSGL4, residues 347–754 a.a. (*C*-terminus); MBY0113016.1, residues 379–774 a.a. (*C*-terminus). (A) Identity matrix of the SGL superfamily. Amino acid sequence identities are shown as percentages (%) in the matrix. These values are colored as a heatmap; the maximum value, cyan; 20%, white; 0%, red. (B) RMSD matrix. The RMSD values are in angstroms. These values were evaluated using the DALI server [27]. PDB and chain IDs are shown beside the protein names. AF2 indicates that AlphaFold2 was used to predict the structures of SkSGL and PgSGL4.

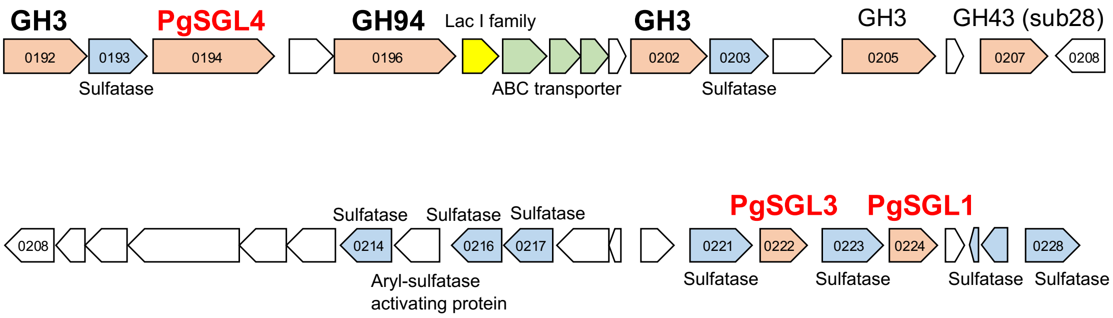

**Figure S3. A gene cluster of genes encoding PgSGL1, PgSGL3, and PgSGL4 from *P. gaetbulicola***
The directions of genes are represented as arrows. The arrow boxes for genes registered in CAZy and SulfAtlas databases [13,14,28,29] are colored light red and light blue, respectively. The arrow boxes for putative ABC transporter genes and a putative LacI family transcriptional regulator gene are colored light green and yellow, respectively. KEGG locus tags with the “H744_1c”, common letters in the loci omitted are shown in the arrow boxes for genes registered in CAZy or SulfAtlas databases. H744_1c0208 is duplicated to show that the genes in the first and the second lines are shown in the same scale. The target proteins in this study are labelled in bold red letters. Proteins with biochemical functions confidently presumed are labelled in bold black letters.

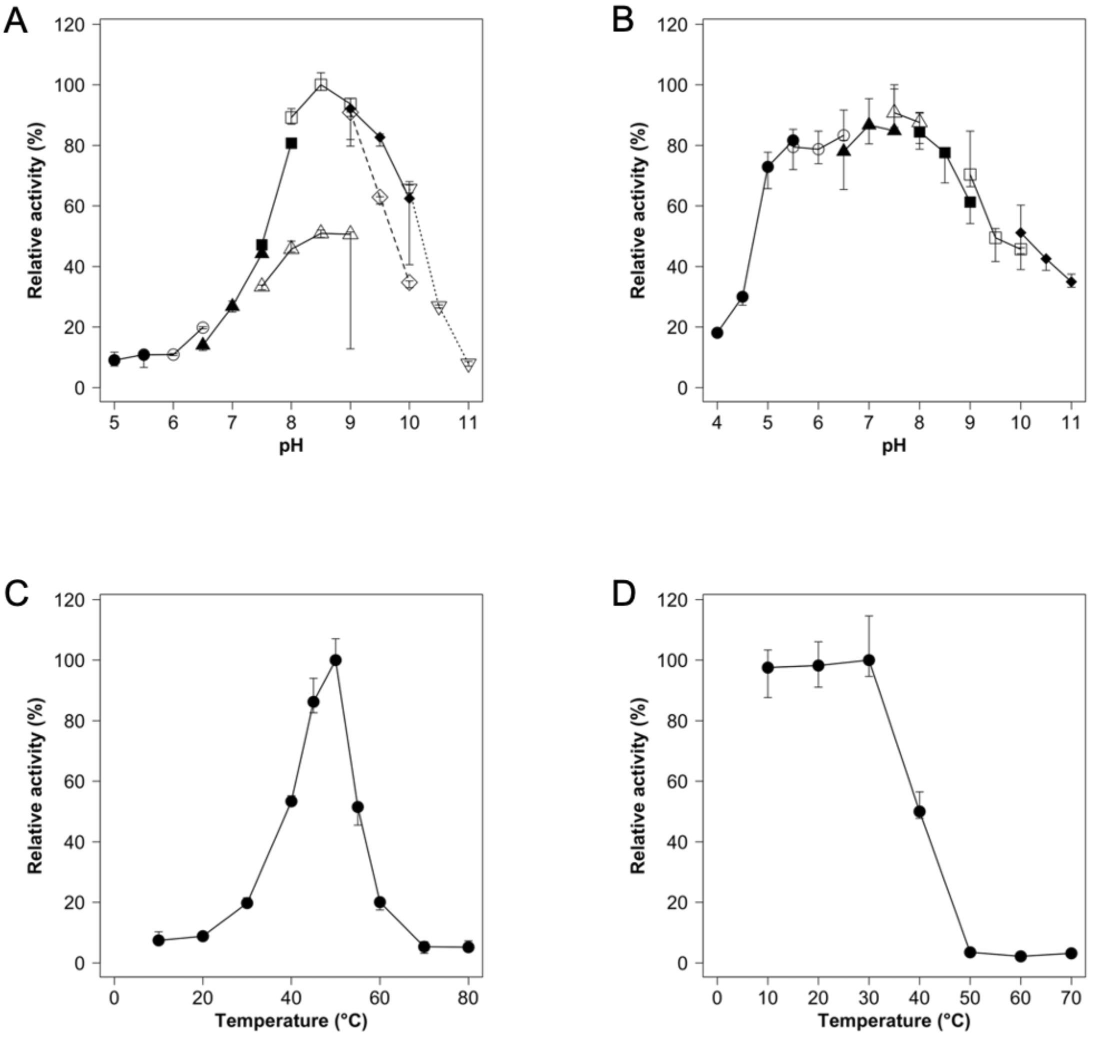

**Figure S4. pH and temperature profiles of PgSGL1 (Group 1)**

The median values of triplicate experiments are plotted as symbols and the other data are shown as error bars. (A, B) investigation of optimal pH (A) and stability (B). Symbols used for optimal pH are closed circles (sodium acetate, pH 5.0–5.5), open circles (MES-NaOH, pH 5.5–6.5), closed triangles (MOPS-NaOH, pH 6.5–7.5), open triangles (Tris-HCl, pH 7.5–9.0), closed squares (HEPES-NaOH, pH 7.5–8.0), open squares (bicine-NaOH, pH 8.0–9.0), closed diamonds (glycine-NaOH, pH 9.0–10.0), open diamonds (CHES-NaOH, pH 9.0–10.0), and open inverted triangles (CAPS-NaOH, pH 10.0–11.0). The counter-ions of the buffers are omitted hereafter. Symbols used for pH stability are closed circles (sodium acetate, pH 4.0–5.5), open circles (MES, pH 5.5–6.5), closed triangles (MOPS, pH 6.5–7.5), open triangles (HEPES, pH 7.5–8.0), closed squares (bicine, pH 8.0–9.0), open squares (glycine, pH 9.0–10.0), and closed diamonds (CAPS, pH 10.0–11.0). (C, D) Optimal temperature (C) and stability (D). Closed circles indicate the median values.

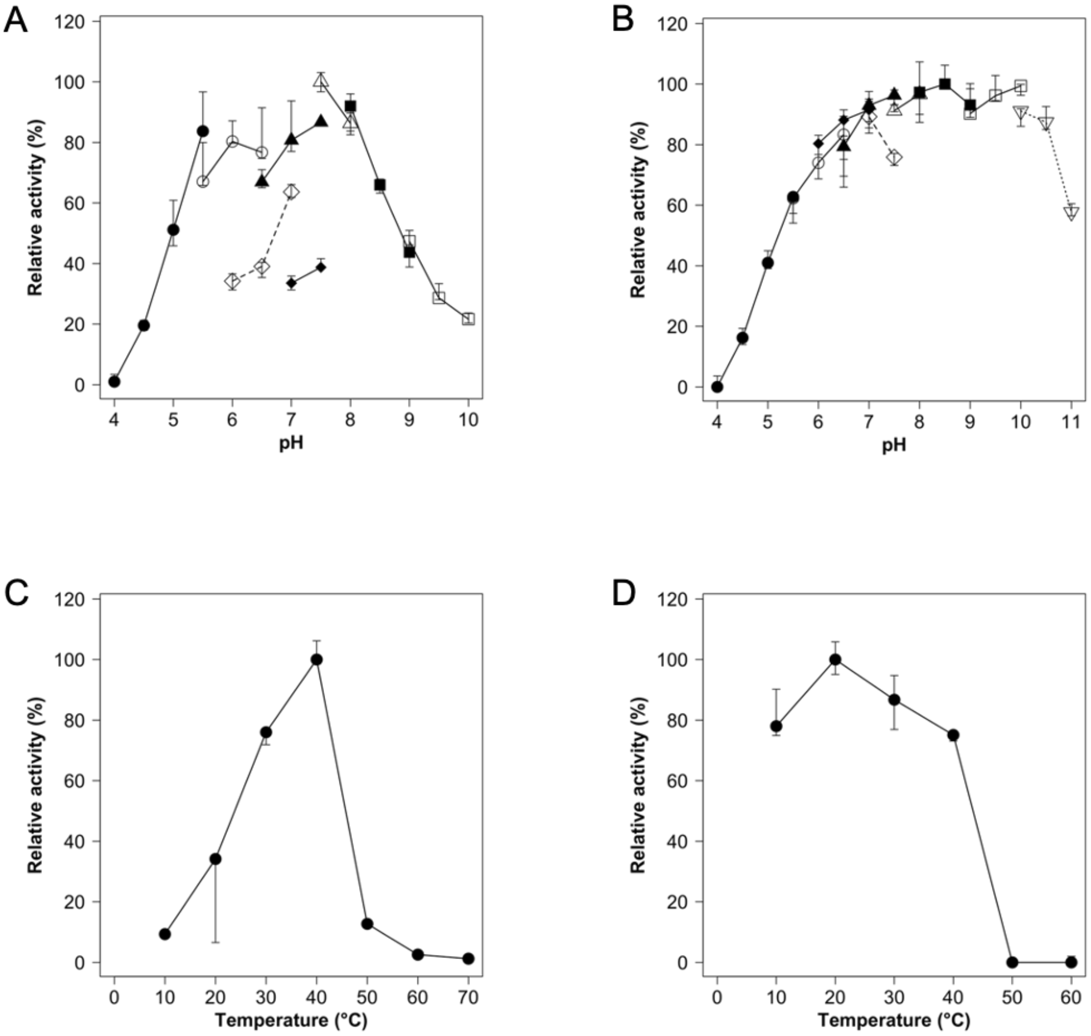

**Figure S5. pH and temperature profiles of PgSGL2 (Group 1)**

The median values of triplicate experiments are plotted as symbols and the other data are shown as error bars. (A, B) investigation of optimal pH (A) and stability (B). Symbols used are closed circles (sodium acetate, pH 4.0–5.5), open circles (MES, pH 5.5–6.5), closed triangles (MOPS, pH 6.5–7.5), open triangles (HEPES, pH 7.5–8.0), closed squares (bicine, pH 8.0–9.0), open squares (glycine, pH 9.0–10.0), closed diamonds (bis-Tris, pH 7.0–7.5), open diamonds (bis-tris propane-HCl, pH 6.0–7.0), and open inversed triangles (CAPS, pH 10.0–11.0). (C, D) Optimal temperature (C) and stability (D). Closed circles indicate the median values.

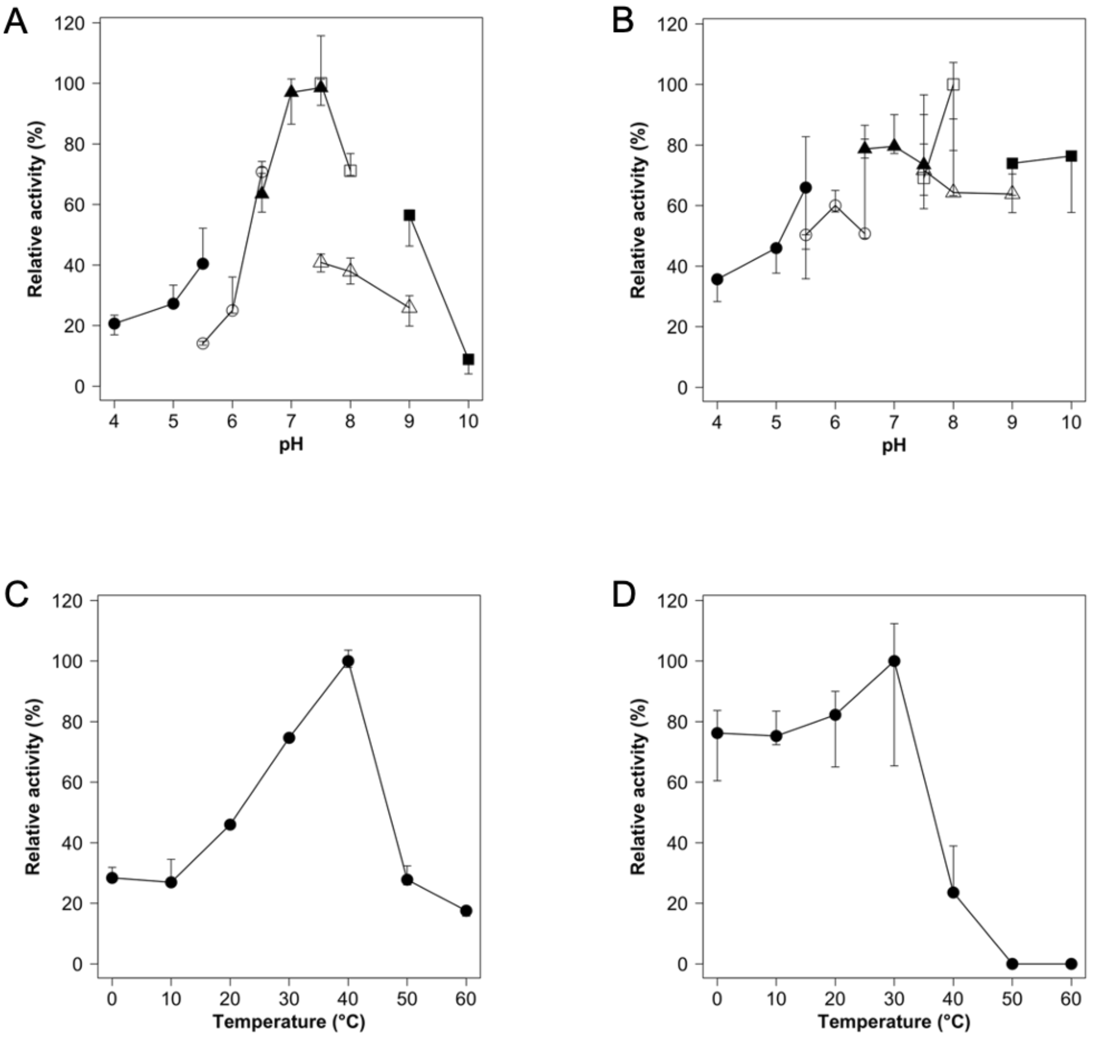

**Figure S6. pH and temperature profiles of EeSGL1 (Group 1)**

The median values of triplicate experiments are plotted as symbols and the other data are shown as error bars. (A, B) Investigation of optimal pH (A) and stability (B). Symbols used are closed circles (sodium acetate, pH 4.0–5.5), open circles (MES, pH 5.5–6.5), closed triangles (MOPS, pH 6.5–7.5), open triangles (Tris, pH 7.5–9.0), closed squares (glycine, pH 9.0–10.0), and open squares (HEPES, pH 7.5–8.0). (C, D) Optimal temperature (C) and stability (D). Closed circles indicate the median values.

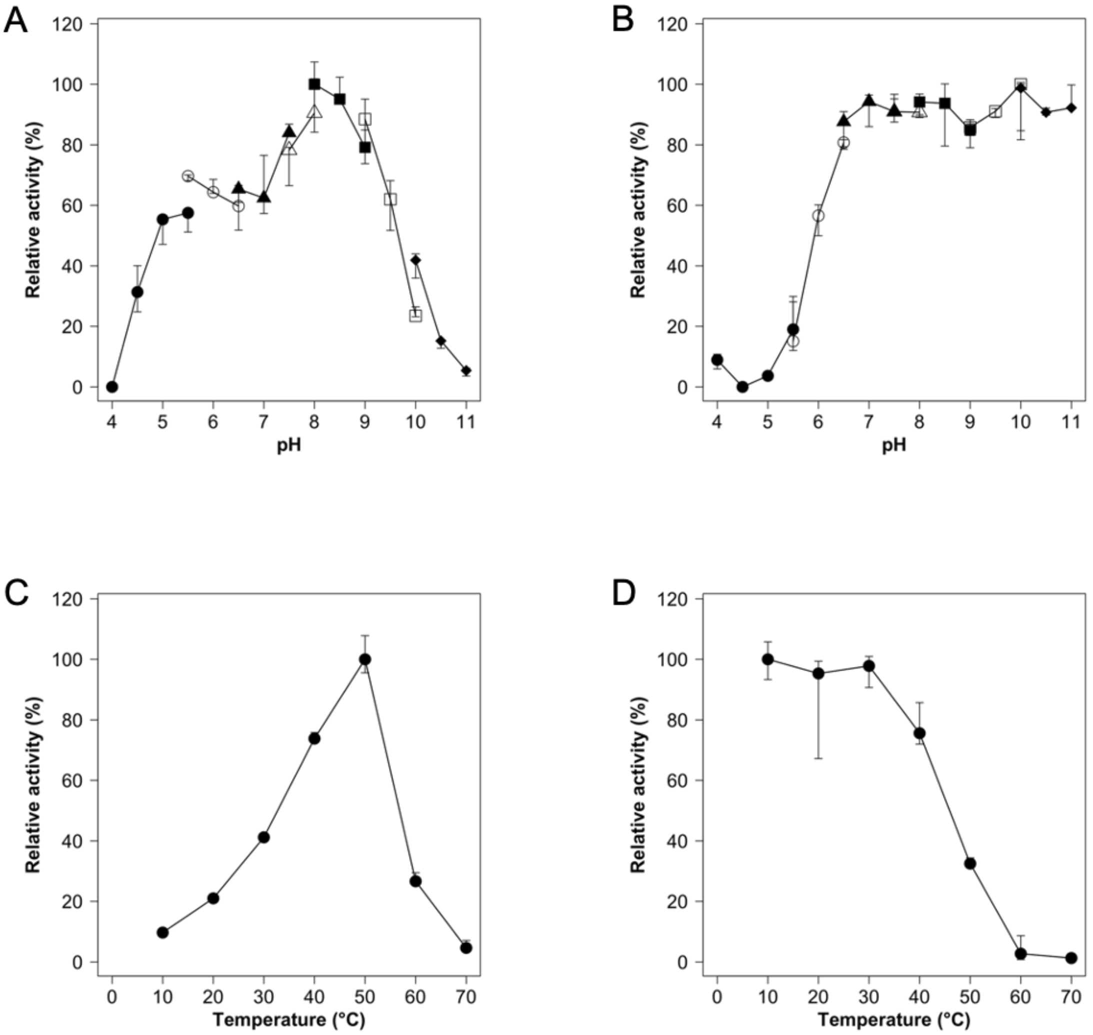

**Figure S7. pH and temperature profiles of SkSGL (Group 2)**

The median values of triplicate experiments are plotted as symbols and the other data are shown as error bars. (A, B) Investigation of optimal pH (A) and stability (B). Symbols used are closed circles (sodium acetate, pH 4.0–5.5), open circles (MES, pH 5.5–6.5), closed triangles (MOPS, pH 6.5–7.5), open triangles (HEPES, pH 7.5–8.0), closed squares (bicine, pH 8.0–9.0), open squares (CHES, pH 9.0–10.0), and closed diamonds (CAPS, pH 10.0–11.0). (C, D) Optimal temperature (C) and stability (D). Closed circles indicate the median values.

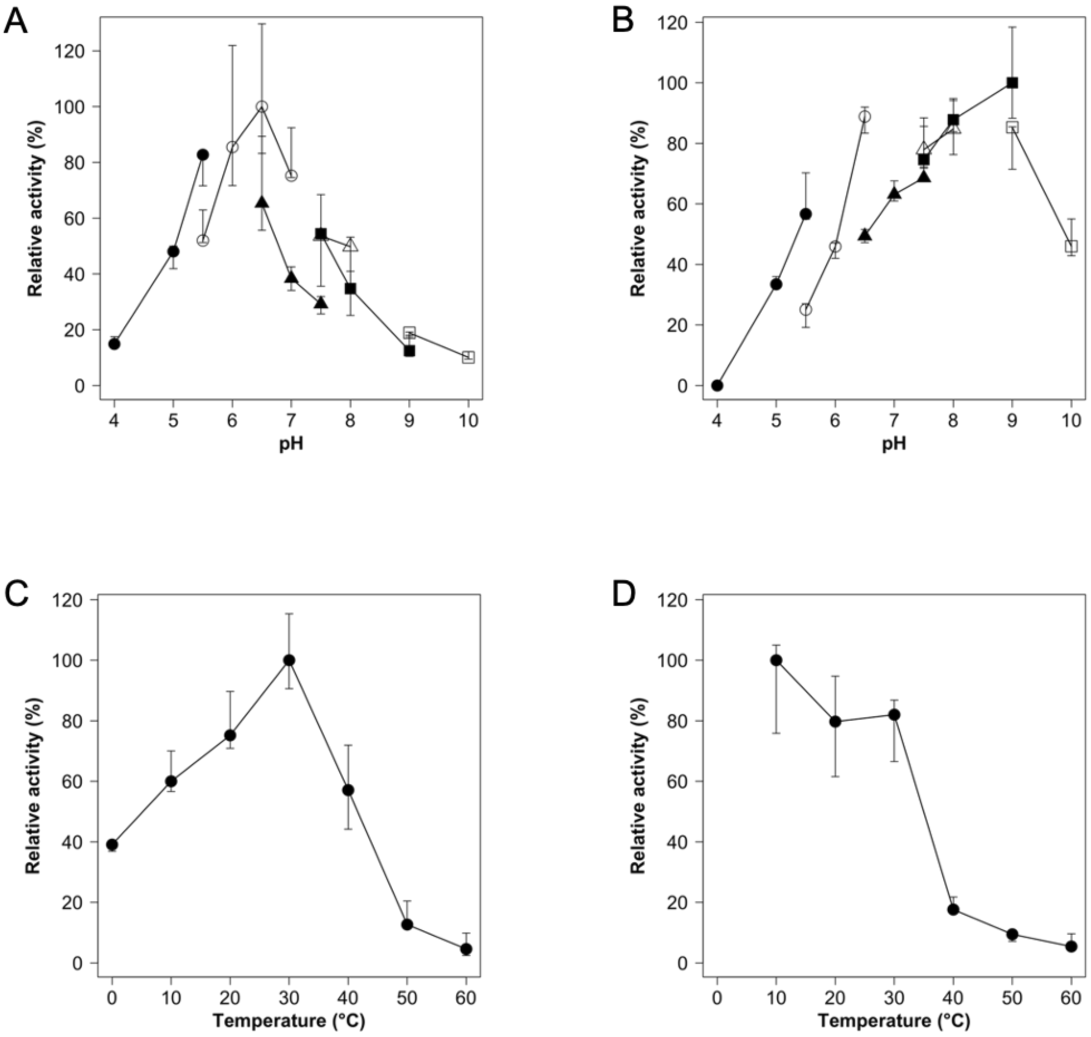

**Figure S8. pH and temperature profiles of PgSGL3 (Group 3)**

The median values of triplicate experiments are plotted as symbols and the other data are shown as error bars. (A, B) Investigation of the optimal pH (A) and stability (B). Symbols used are closed circles (sodium acetate, pH 4.0–5.5), open circles (sodium cacodylate, pH 5.5–7.0), closed triangles (MOPS, pH 6.5–7.5), open triangles (HEPES, pH 7.5–8.0), closed squares (Tris, pH 7.5–9.0), and open squares (glycine, pH 9.0–10.0). (C, D) Optimal temperature (C) and stability (D). Closed circles indicate the median values.

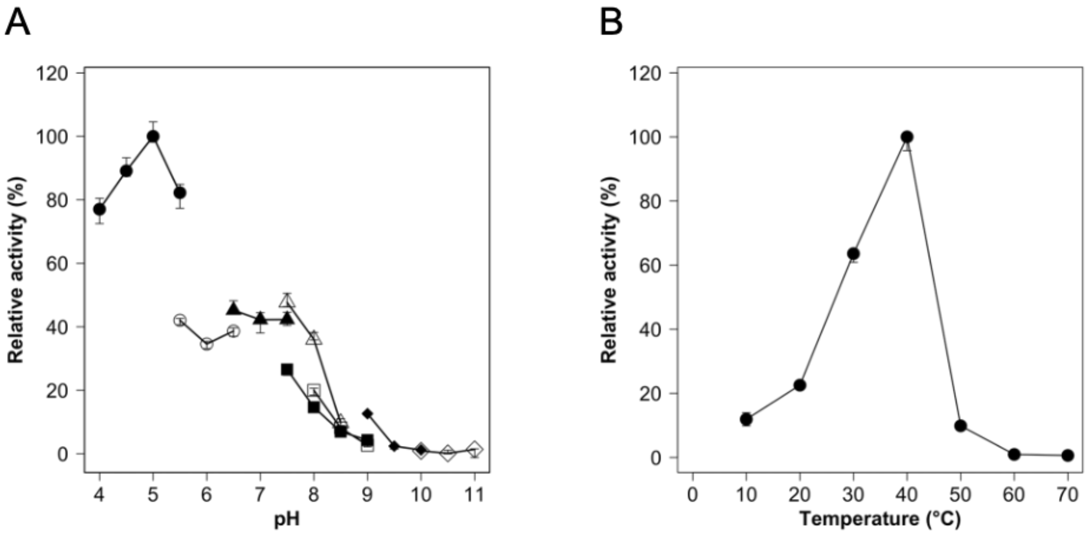

**Figure S9. Optimal pH and temperature of XcSGL (GH144)**

The median values of triplicate experiments are plotted as symbols and the other data are shown as error bars. (A) Investigation of the optimal pH. Symbols used are closed circles (sodium acetate, pH 4.0–5.5), open circles (MES, pH 5.5–6.5), closed triangles (MOPS, pH 6.5–7.5), open triangles (HEPES, pH 7.5–8.5), closed squares (Tris, pH 7.5–9.0), open squares (bicine, pH 8.0–9.0), closed diamonds (glycine, pH 9.0–10.0), and open diamonds (CAPS, pH 10.0–11.0). (B) Investigation of the optimal temperature. Closed circles indicate the median values.

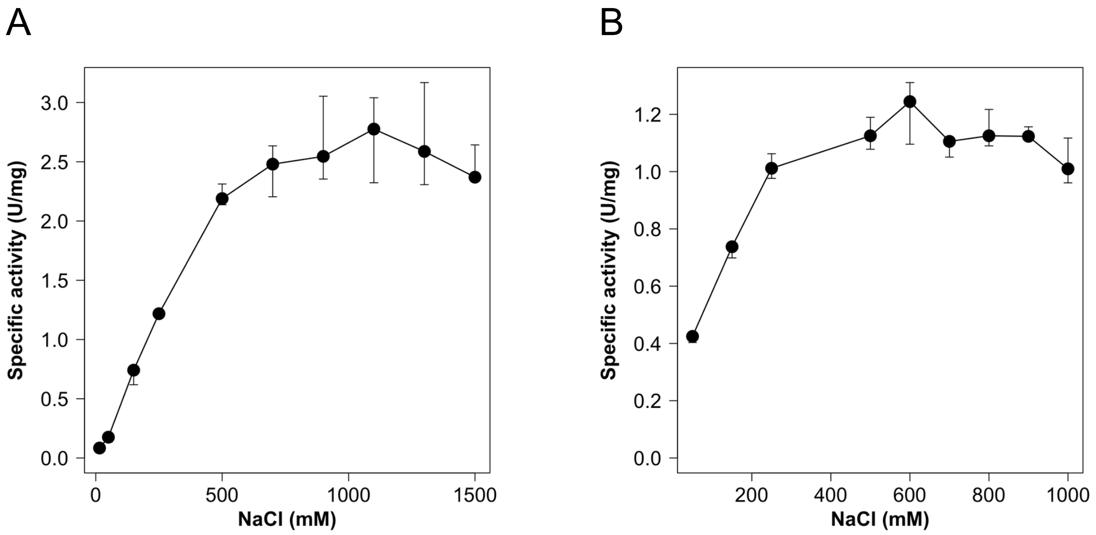

**Figure S10. Effect of NaCl on the hydrolytic activity of PgSGL2**

Hydrolytic activity (A) and stability (B) of PgSGL2 in the presence of various concentrations of NaCl. Medians in the triplicate experiments are plotted as closed circles and the other data are shown as error bars.

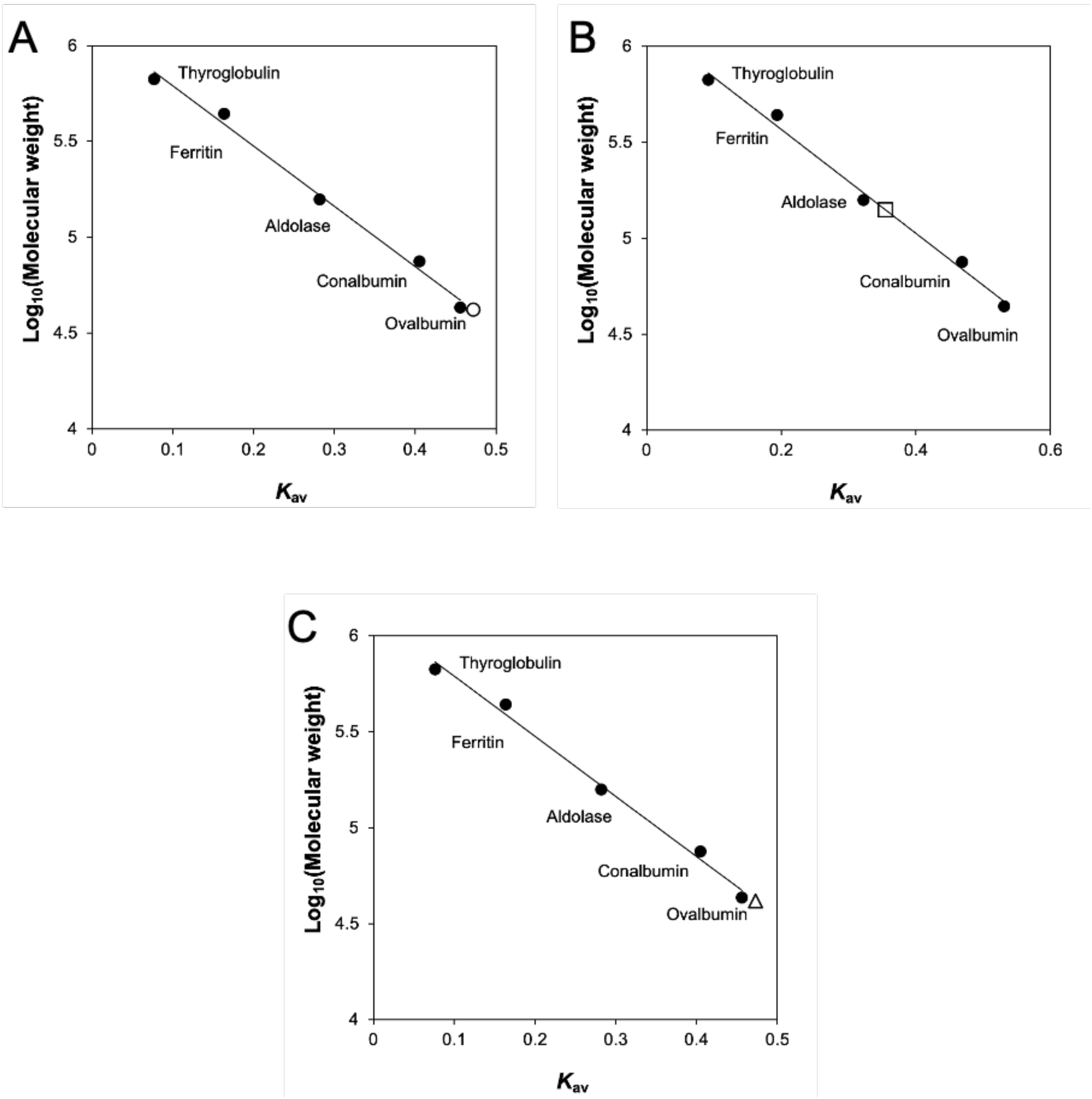

**Figure S11. Size-exclusion chromatography analysis**

(A), EeSGL1 (Group 1); (B), SkSGL (Group2); and (C), PgSGL3 (Group 3). Protein standard markers are shown as closed circles. EeSGL1, SkSGL, and PgSGL3 are shown as open circle, square, and triangle, respectively.

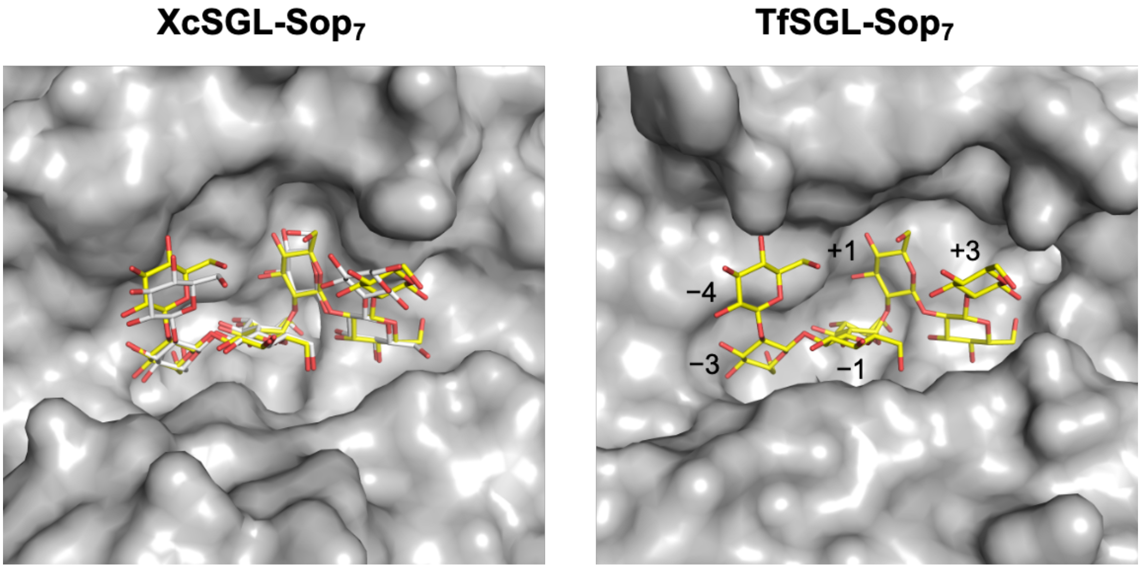

**Figure S12. Comparison of substrate pockets between XcSGL-Sop$_7$ and TfSGL-Sop$_7$ complexes.**

E239Q mutant of XcSGL and E262Q mutant of TfSGL are used for preparation. XcSGL and TfSGL are shown as gray surfaces. Sop$_7$ molecules in XcSGL and TfSGL are shown as white and yellow sticks, respectively.

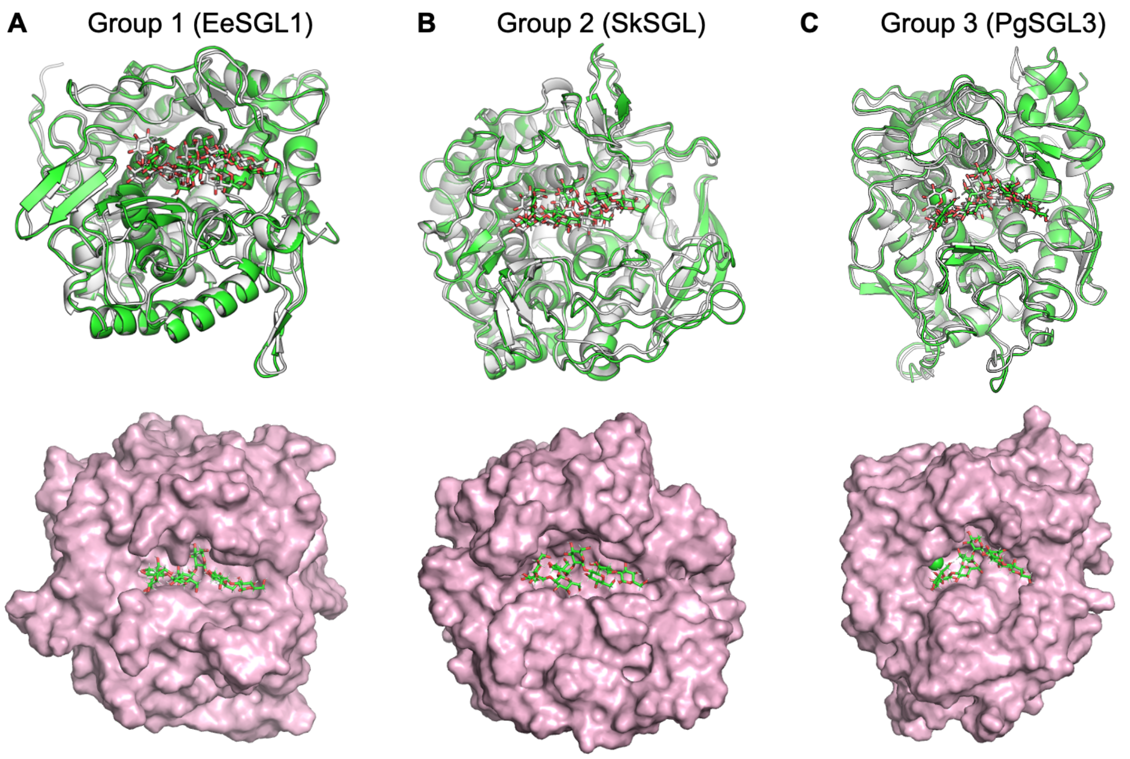

**Figure. S13. Structures of the complexes obtained by MD simulations**

(A–C) Structures before and after MD simulations are colored white and green, respectively. (top) Proteins and substrates are shown as cartoons and sticks, respectively. (bottom) Final structures in the MD simulations. The proteins and substrates are shown with pale pink surfaces and as green sticks, respectively. (C) A chloride ion is shown as a sphere.

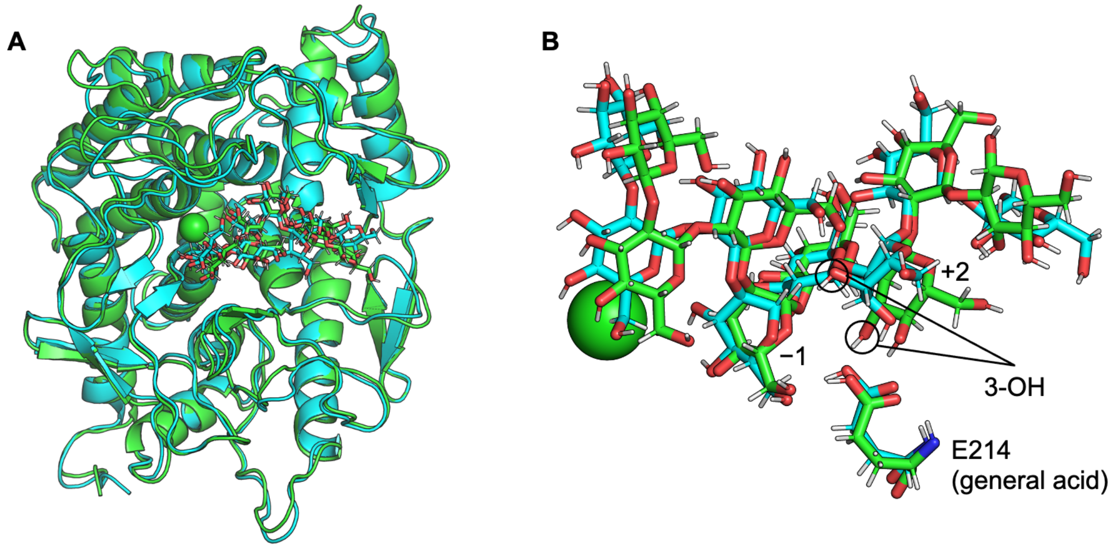

**Figure S14. Structures of PgSGL3 complexes in the presence and absence of a chloride ion**

The structures with and without a chloride ion in the catalytic pocket are shown in cyan and green, respectively. $Sop_8$ molecules are shown as sticks. (A) Overall structures of the complexes of PgSGL3 with $Sop_8$ obtained by MD simulations. (B) Enlarged view of the $Sop_8$ molecules, E214 residues, and chloride ions in the catalytic pocket. Subsite numbers are shown at the substrates.

```
                         α1                                β1
CpSGL_GH144                                 T.....T...T        TT               TT  TT
CpSGL_GH144    25  S.....ADSIFNIVEEQTFQYFWD...GAEP.....V...SGMARERYHVD..........GNYPENDMN
XcSGL_GH144    57  ......LPPLFSDIERRTFQFFWD...TTNE.....L...NGLSPDRFP................S....RP
EeSGL1         38  .......PADFREELQTNLNF.FLDGKGVDA.....D...TLVPYDTIWVK....D.N.K.AEY....AY
SkSGL          36  ....PEQTAVLERYVTDTYASLAA...MTDD.....T...TGLPADNIGG.......DLDPASA....SA
PgSGL3         37  D....Q..EIIDSLYRGGYAYWQQ...LRNE.........NGTYEDKLFL.........NGD.R....SY
PgSGL4        347  D.....FDAWMRDELAQYVAAIKG...LTASGMSGNYASPEGFLHRKLEH..T........NPN....TS
TiCGS_GH189  1013  ...SQEEMEELRLVARKTWRYFED...FVTE.....G.Q.NYLPPDNFQEDPPN........GV....AE
TfSGL_GH162    37  QVLQ.NPSKFINDVLFWEGKF.HQNNISYNS.....G...NGMSYDGTNIDW..VTGE.G.TVK....HP

                   β2         α2                                α3
CpSGL_GH144
CpSGL_GH144    69  VVTSGGSGFGVMALLVGI..ER..G..YI.....S.REQGLERLMKIVSFLEK......A..........
XcSGL_GH144    92  FASIASVGFALTAYPIGI..EN..G..WV.....S.RNQAIDRTLTTLKFFRD......A..P.MGPQRT
EeSGL1         81  YTNTTEIALYLNILVEAE.K....A.........GN..QKALTRIQEVLTTLEE......A...PK.....
SkSGL          80  YTSPTNIGGYLWSTVMAR..DL..G..IV.....T.VDEAYDLMSTTLATVEG......L...D......
PgSGL3         75  VGSIANSGMGLIALTIGH..AN..G..WE.....P...EAEQLALVTLRKLAGRDPN.FA...V......
PgSGL4        395  PIAD.NVGWALYLLMVAD..QIE.H...D.....P...EIEQYIELLIKRHAGLHEDGKGG..V......
TiCGS_GH189  1058  RTSPTNIGLYLVSVIGAR..DL..G..YI.....T.TTEMVERIKKTLDTIEK......M...E......
TfSGL_GH162    89  FSAASKESLQVMLYAHAIAGSADAARFLSPNNPSAAPGIAASIMDTKLQTYLR......FNETYPG....
                      ★

                                β3                           β4      α4
CpSGL_GH144                TT                               TT
CpSGL_GH144   111  ..DRF..HGAWPHWLYG..ETGKVKP.........FGQKD.NGGDLVETSFMIQGLLCVRQYFAN.G...
XcSGL_GH144   141  GKAGY..KGFYYHFLDM..QQGNRYD.........S..W.V.E.LSSVDTALLMMGVLFTQSYYD..G...
EeSGL1        122  ....F..KGLFYWPYDIKGGELKPGK.............G.EIAPAVDNGNLAFSLAAVAGAYLN.S...
SkSGL         123  ...RHEPSGMFYNWYDP..ATGERVRNWPGDGVVV..D..Q.FLSSVDNGWLAAALRVVAEAEP......
PgSGL3        121  ..PQN.ATNTFIHFYNT..KTGEAV..........G...D..DWSPVDSAIMIYGALFVKNYFS..E...
PgSGL4        442  ...KS.VDGHFVRNYN.TD.GSI.NA.......GN.....P.QYQVYISMKFLPAAIKAAEMYP......
TiCGS_GH189  1101  ...KW..NGHLYNWYNT..KTLEPL..........R..P..Y.YVSTVDSGNLVGYLITVKEAIGEFLNKP
TfSGL_GH162   149  ....F..GGFLP.WFTSSSQDLTPTW...........D.WNNRVPGLDNGELLWAVYAFIQAAEN.T...
                                                                   ★

CpSGL_GH144
CpSGL_GH144        ......................................................................
XcSGL_GH144        ......................................................................
EeSGL1             ......................................................................
SkSGL              ......................................................................
PgSGL3             ......................................................................
PgSGL4             ......................................................................
TiCGS_GH189  1150  LIDIELAKGLKDTIKMLNVKGITEDIFTTILSKTTLVPSEWEAFLNKIREKLLSSQDDLENIKRLKNIII
TfSGL_GH162        ......................................................................

CpSGL_GH144
CpSGL_GH144        ......................................................................
XcSGL_GH144        ......................................................................
EeSGL1             ......................................................................
SkSGL              ......................................................................
PgSGL3             ......................................................................
PgSGL4             ......................................................................
TiCGS_GH189  1220  ALKGEMKEFLVWTEFDESEKEQEIFKRYKEVFEEHSSPKELEKVYKNYLLEIEEVFKKATEEEKALLKSQ
TfSGL_GH162        ......................................................................

                                      α5              η1
CpSGL_GH144
CpSGL_GH144   161  ..............NEQEKALAARIDQLWKAV...EFSW.YRN....GK................NVLY
XcSGL_GH144   189  .............DDPREKEIRQIADTLYKRV...DWRW.LQQ..R..A.................PLIS
EeSGL1        168  .............TDPVKQSIISRIDQMLKAQ.IPGWLS.LYD....KDR.................GLLW
SkSGL         177  .................R..LADEATAVYDDM...NFG.AFYN....ADA.LPDRGL.........GLLR
PgSGL3        166  .................NEEIAELADFLYRNT...DLTQ.YI...A...D.......VR...TG...RIY
PgSGL4        486  ..............D..NQNIASYAKYLQQVF.Q.RAGDVVRAEQRI..TW.....DSDDFGPVR.....
TiCGS_GH189  1290  KDKVAQALEKIKKL.EAEIENIKSTIENLVEKT...EFR.HLYD....EKR.................QLFS
TfSGL_GH162   196  .............SNKSFIDLAKKWQTWMDYTKTT.AAHIFYQ....GE.................GKVC

                   β5                                                  α6
CpSGL_GH144                                                                         TTT
CpSGL_GH144   192  WHWSPNY..........................KWQM.NFP..VTG.YNECLIMYILAAAS..PTH.....
XcSGL_GH144   221  MGWFPES..........................GF.I.DHD..WMG.YNEAMMLYILALGS..PTH.....
EeSGL1        203  GGWQNG...........................EL.I.EYH..VDRKANESRLAALWAPLI....TKHLGA
SkSGL         210  GGFWDEVQTEGSVPGDYLG......T.G.TEVFY.T.GHH..YDTQVSETRIATYLGIAE.......G....
PgSGL3        196  LAQHTD...........................GTF.KK.YR..TKA.FNEYMLVAGIANQQAKDL...D..
PgSGL4        526  ...................................LNRLM.S..NETWLYGDIAAAQD....P.....
TiCGS_GH189  1336  IGYNVEE..........................EKL.T.KSY..YDLLASEARQASFIAIAK.....K....
TfSGL_GH162   231  AVTDIK............NQSLPVYHPEQTYAC.EGTSY..LNDP.YEGELFTWWLQFF....G.....
                                                                ▲
```

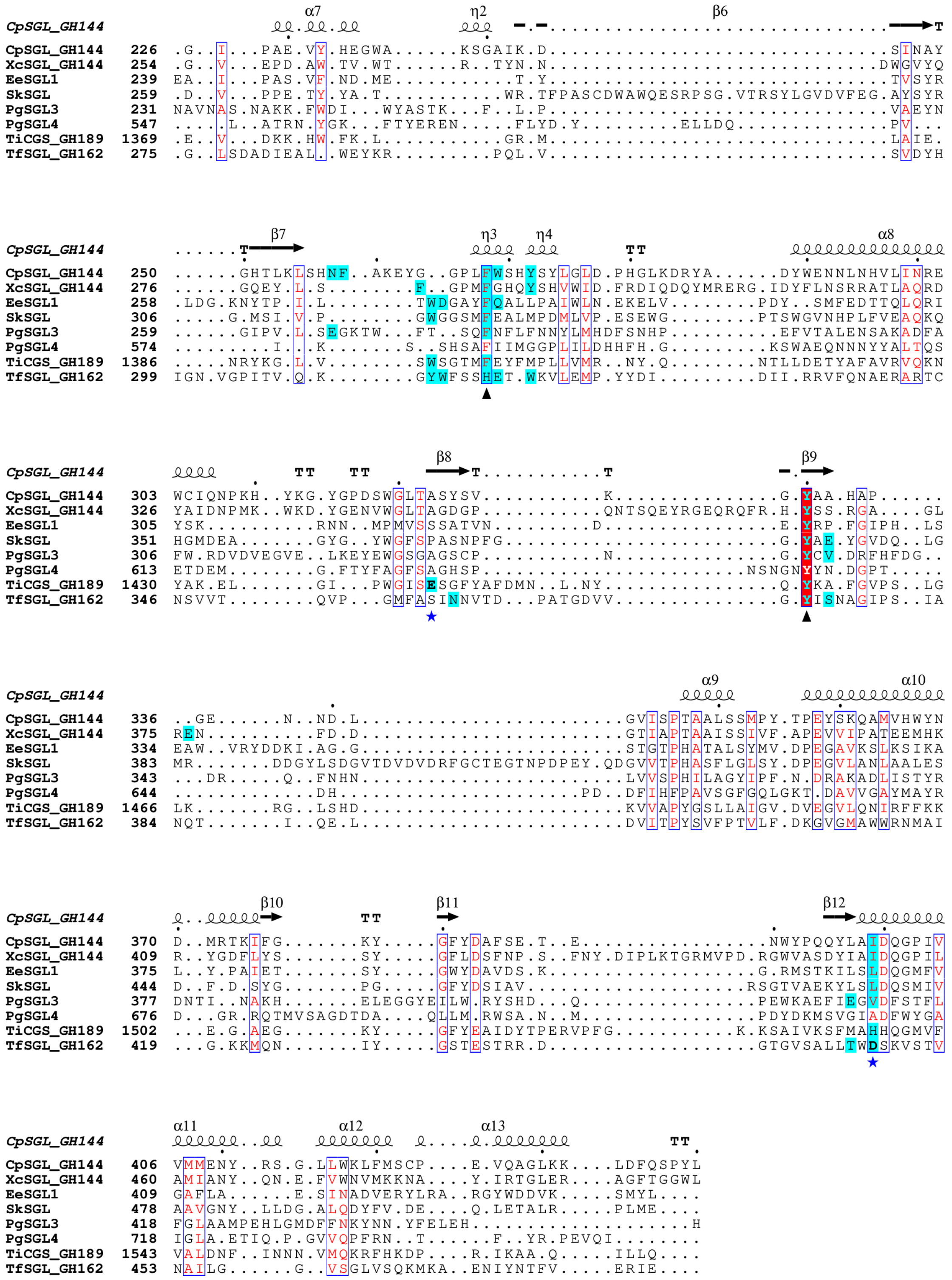

**Fig. S15. The entire multiple sequence alignment of the SGL-superfamily proteins**

A version of the entire multiple alignment shown in Fig. 7. Residues in bold letters labelled with a blue star are a general base catalyst for TfSGL, a nucleophile for TiCGS$_{Tg}$, and candidate general base catalysts for the other proteins. The residues that define the SGL-superfamily proteins are labelled with black triangles. Residues within 4 Å of the superimposed $Sop_7$ molecule from the XcSGL (E239Q)-$Sop_7$ complex are colored cyan.

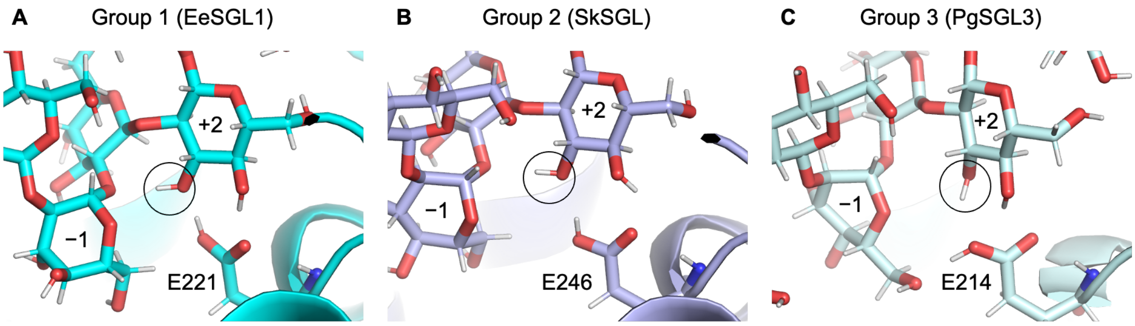

**Figure S16. Substrate recognition by the catalytic residues**

Substrates (Sop$_8$ molecules) and proteins (A, EeSGL1; B, SkSGL; and C, PgSGL3). EeSGL1, SkSGL, and PgSGL3 are colored the same as the corresponding proteins in Fig. 5 (cyan, light purple, and pale cyan, respectively). The complex structures are the final structures in the MD simulations. 3-Hydroxy groups are indicated by black circles. The candidates for general acids are shown as sticks. Subsite numbers are shown at the substrates.

**A** PgSGL1 HisTrap$^{TM}$ crude

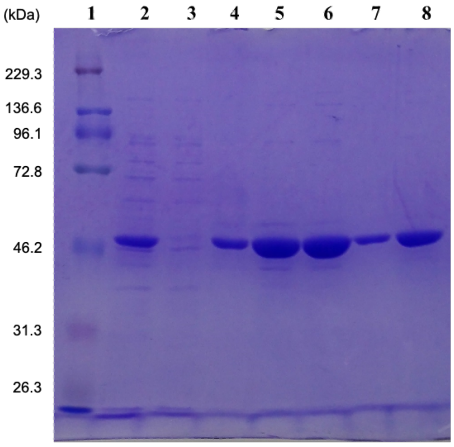

1, Protein marker
2, Crude extract
3, Flow through
4, Fraction 4
5, Fraction 5
6, Fraction 6
7, Fraction 8
8, Fraction 7

**B** PgSGL2 HisTrap$^{TM}$ crude

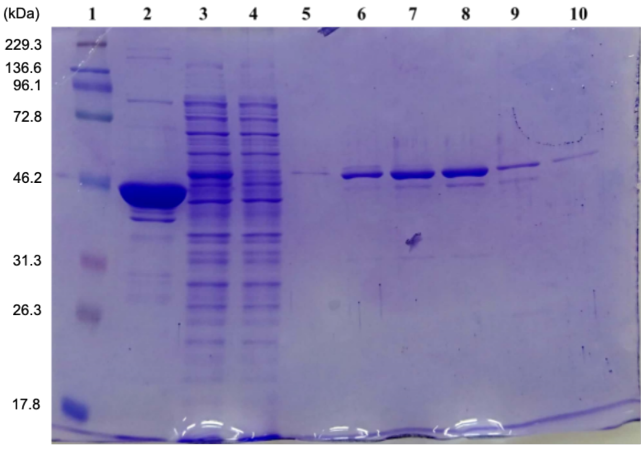

1, Protein marker
2, (not related to this study)
3, Crude extract
4, Flow through
5, Fraction 3
6, Fraction 4
7, Fraction 5
8, Fraction 6
9, Fraction 7
10, Fraction 8

**C** EeSGL1 HisTrap$^{TM}$ crude

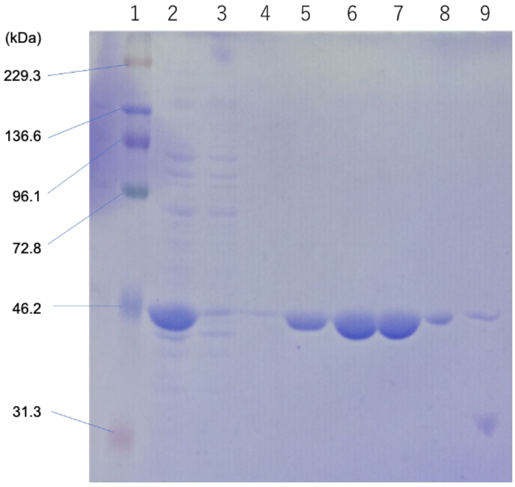

1, Protein marker
2, Crude extract
3, Flow through
4, Fraction 4
5, Fraction 5
6, Fraction 6
7, Fraction 7
8, Fraction 8
9, Fraction 9

**D** PgSGL3 HisTrap$^{TM}$ crude

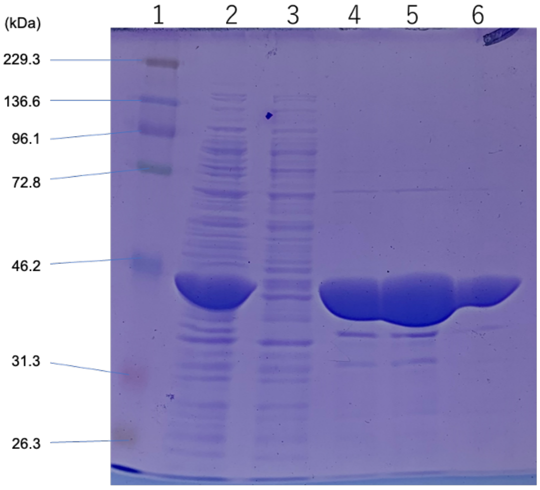

1, Protein marker
2, Crude extract
3, Flow through
4, Fraction 5
5, Fraction 6
6, Fraction 7

**E** XcSGL HisTrap™ crude

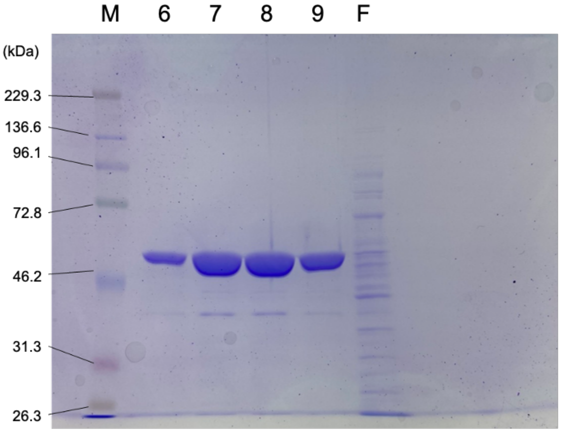

M, Protein marker
6, Fraction 6
7, Fraction 7
8, Fraction 8
9, Fraction 9
F, Flow through

**F** SkSGLn HisTrap™ crude

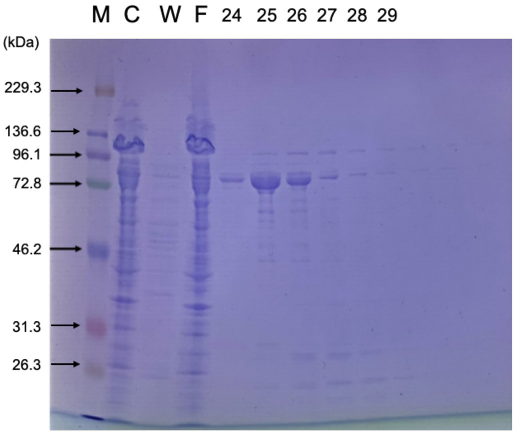

M, Protein marker
C, Crude extract
F, Flow through
24–29, Fraction numbers

**G** SkSGLc

HiTrap™ DEAE FF

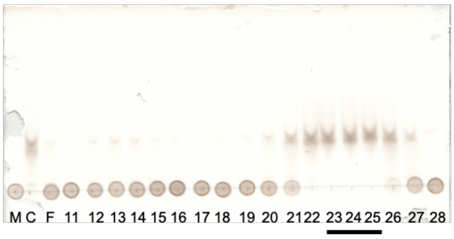

HiTrap™ Butyl HP

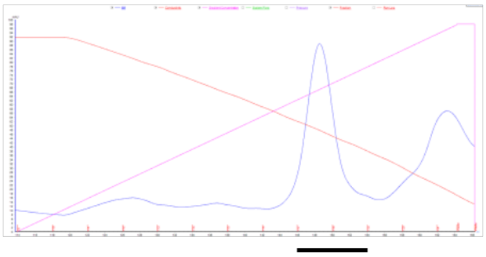

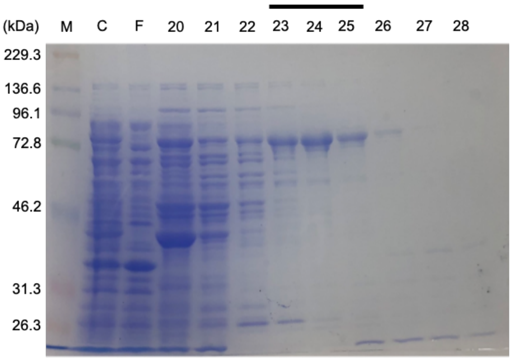

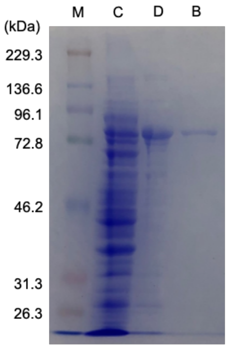

M, Protein marker; C, Crude extract; F, Flow through: D, mixture of DEAE fraction 23–25; B, SkSGL used for assay after purification byHiTrap-Butyl HP; Numbers, Fraction numbers. Bars, Collected fractions

**Figure S17. Purification of SGL-superfamily enzymes**

The enzymes and columns used for purification are shown above the SDS-PAGE gels, TLC plate, and chromatography chart. Descriptions of the lanes for SDS-PAGE are shown below the SDS-PAGE gels. Molecular weights of the protein markers are shown beside the SDS-PAGE gels. (G) (top left) TLC analysis of fractions. The reactions were performed at 30 °C for 3 h in solutions (10 μl) containing 0.5% β-1,2-glucan (DP121), 50 mM HEPES (pH 8.0), and 1 μl of each fraction. Each reaction solution (1 μl) and a marker were spotted on the TLC plate. Lane M, 0.5% β-1,2-glucan (DP121) as a marker. (top right) Chromatography chart. The blue, red, and magenta lines represent the UV absorbance, conductivity, and percentage of elution buffer, respectively, according to ÄKTA™ prime (Cytiva).

https://doi.org/10.1093/nar/gkab1045.
